# The inflammatory and tumor suppressor SAMD9L acts through a Schlafen-like box to restrict HIV and inhibit cell translation in SAAD/ATXPC

**DOI:** 10.1101/2023.01.19.524725

**Authors:** Alexandre Legrand, Clara Dahoui, Clément De La Myre Mory, Clara Loyer, Kodie Noy, Laura Guiguettaz, Margaux Pillon, Mégane Wcislo, Laurent Guéguen, Andrea Cimarelli, Mathieu Mateo, Francesca Fiorini, Emiliano Ricci, Lucie Etienne

**Author notes:** Correspondence: Lucie Etienne.

## Abstract

Sterile alpha motif domain-containing proteins 9 and 9L (SAMD9/9L) are associated with life-threatening genetic diseases and are restriction factors of poxviruses. Yet, their cellular function and the extent of their antiviral role are poorly known. Here, we found that interferon-stimulated SAMD9L, and not SAMD9, restricts HIV-1 replication at the translation step, with a strong inhibition of Transmitted/Founder HIV-1 patient strains. More broadly, SAMD9L restricts primate lentiviruses, but not another retrovirus (MLV) or two ssRNA viruses (MOPV, VSV). Using structural modeling and mutagenesis of SAMD9L, we identified a Schlafen(SLFN)-like active site necessary for HIV-1 restriction. By testing a germline gain-of-function variant from patients with SAMD9L-associated autoinflammatory disease (SAAD) and ataxia-pancytopenia (ATXPC), we determined that SAMD9L cellular and pathogenic functions also depend on the SLFN-like active site. Finally, we propose a model in which SAMD9L translational repression could be dependent on codon-usage, linking its cellular function and the virus-specific innate immunity. The identification of another Achille’s heel of HIV, as well as the inflammatory SAMD9L effector and auto-regulatory determinants, provide novel avenues against infectious and genetic diseases.

**Significance statement:** This study identifies SAMD9L as a potent HIV-1 antiviral factor from the interferon immunity and deciphers the host determinants underlying SAMD9L translational repression. The characterization of SAMD9L activity and determinants is also of medical importance for patients with rare genetic diseases bearing deleterious mutations in SAMD9L or with specific cancers. We demonstrate that a pathogenic SAMD9L patient’s variant is inactivated by the mutation of an identified active site in a SLFN-like box, resulting in an abolished translational shutdown. Furthermore, we describe SAMD9L, but not SAMD9, as an antiviral factor of HIV and lentiviruses, through a translational repression mediated by the SLFN-like box and potentially dependent on codon usage. These findings may have implications to better fight against HIV/AIDS as well as SAAD/ATXPC.

**Key findings:**

- SAMD9L, but not SAMD9, restricts HIV-1, including Transmitted/Founder patient strains.
- SAMD9L broadly restricts primate lentiviruses, but not the retrovirus MLV, nor two ssRNA viruses, the Rhabdovirus VSV and the Arenavirus MOPV.
- SAMD9L inhibits viral and cellular translation through an essential E198/D243 active site in a SLFN-like box.
- The SAMD9L-associated autoinflammatory disease (SAAD) F886Lfs*11 variant has enhanced HIV translational repression, unveiling an autoregulatory domain of the anti-lentiviral function.

## Introduction

The Sterile Alpha Motif Domain-containing 9 (*SAMD9*) and *SAMD9L* are paralogous genes that encode large proteins of more than 1500 amino acids with a complex architecture containing multiple domains. SAMD9 and SAMD9L are involved in several cellular processes such as cell proliferation and protein translation^1–3^, apoptosis and stress responses^1^, as well as endosomal trafficking^4,5^. Both genes are involved in life-threatening genetic diseases. Genetic variants of SAMD9 are associated with a multisystem disorder, MIRAGE, and SAMD9L genetic variants are responsible for neurological and hematological disorders, such as ataxia-pancytopenia (ATXPC) or SAMD9L-associated autoinflammatory disease (SAAD)^2,6–8^. These syndromes reflect gain-of-function (G-o-F) phenotypes caused by missense mutations or truncated forms. Mechanistically, the SAMD9/9L associated diseases are probably a consequence of accrued translational repression by the G-o-F variants.

SAMD9 and SAMD9L are also restriction factors of poxviruses, acting through virus translational repression. In turn, mammalian poxviruses encode species-specific antagonists, from the C7K superfamily, which are necessary for viral replication and pathogenesis^9–11^. Although double-stranded nucleic acid (dsNA) binding appears an important determinant of anti-poxviral activity and antiproliferative activity of SAMD9/9L G-o-F variants^12^, the effector functions of SAMD9/9L and the extent of their antiviral immune activity remain unknown.

The Human Immunodeficiency Virus Type 1 (HIV-1) is responsible for acquired immunodeficiency syndrome (AIDS) and remains a major public health concern with 38 million people living with the virus in 2021 (UNAIDS). The identification and characterization of cellular proteins that naturally inhibit HIV and related viruses have been key to understanding HIV infection and mammalian antiviral innate immunity^13–15^. Furthermore, several factors at the interface with HIV are also involved in major cellular dysfunctions, such as the HIV restriction factor SAMHD1 also involved in Aicardi-Goutières syndrome (AGS)^16^. Overall, unveiling host factors that impact HIV at different steps of replication and that are possibly involved cellular dysfunctions helps to design new drug targets against viral pathogens and genetic diseases.

Here, by combining protein structure and genetic analyses, with molecular virology and functional characterization, we show that the interferon-induced SAMD9L, and not SAMD9, specifically inhibits HIV and other lentiviruses, through a newly-identified Schlafen-like (SLFN) active site driving SAMD9L cellular and antiviral functions. Our study therefore identifies a novel ISG-encoded protein with antiviral functions against HIV, and further provides a link between SAMD9L functions in antiviral immunity and life-threatening syndromes.

## Results

### SAMD9L, and not SAMD9, acts as an antiviral factor against HIV-1 and primate lentiviruses

SAMD9L has been recently retrieved as a potential HIV modulator in a genome-wide CRISPR-based screen^17^. However, its validation and functional relevance in the context of HIV infection has not been investigated yet. To determine if SAMD9 and SAMD9L restrict HIV-1, we tested the capacity of the ectopically-expressed proteins to impact the infectivity of four replication-competent full-length HIV-1 viruses. We tested two HIV-1 laboratory-adapted strains (LAI and NLADA) and two Transmitted/Founder (T/F) strains (pCH077 and pWITO), which correspond to initial viral variants responsible for a productive infection in HIV-1-infected patients ^18,19^. To do so, we co-transfected 293T cells with or without a SAMD9/9L-encoding plasmid and one of the HIV-1 infectious molecular clones (IMCs), and we titered the HIV-1 infectious yield in the supernatant by infecting the TZM-bl reporter cell line, which encodes for luciferase downstream of an LTR promoter (Fig. 1A). We found that SAMD9L inhibited HIV-1 infectivity, while SAMD9 exerted an opposite, enhancing HIV-1 (Fig. 1B). In these conditions, exogenous SAMD9/9L did not impact cell counts, showing that the anti/pro-viral effects were independent of the overall cell proliferation (Fig. S1). Interestingly, SAMD9L strongly restricted HIV-1 T/F strains – with one-to-two log restriction in infectivity –, whereas HIV laboratory-adapted strains were less impacted (Fig. 1B). Viral doses (i.e. decreasing/increasing amounts of HIV-1 IMCs in the producer cells; Fig. S2A-B) confirmed a differential susceptibility between HIV-1 strains, with SAMD9L strongly restricting T/F pWITO compared to pLAI for example, suggesting some intrinsic HIV-1 strain-specific susceptibility effect to SAMD9L restriction. Furthermore, the inhibition of HIV-1 was dependent on the dose of SAMD9L (Fig S2C) – a typical feature of antiviral factors. Because of the strong antiviral effect of SAMD9L on HIV-1, we concentrated the rest of the study on this immune defense protein.

**Figure 1.**
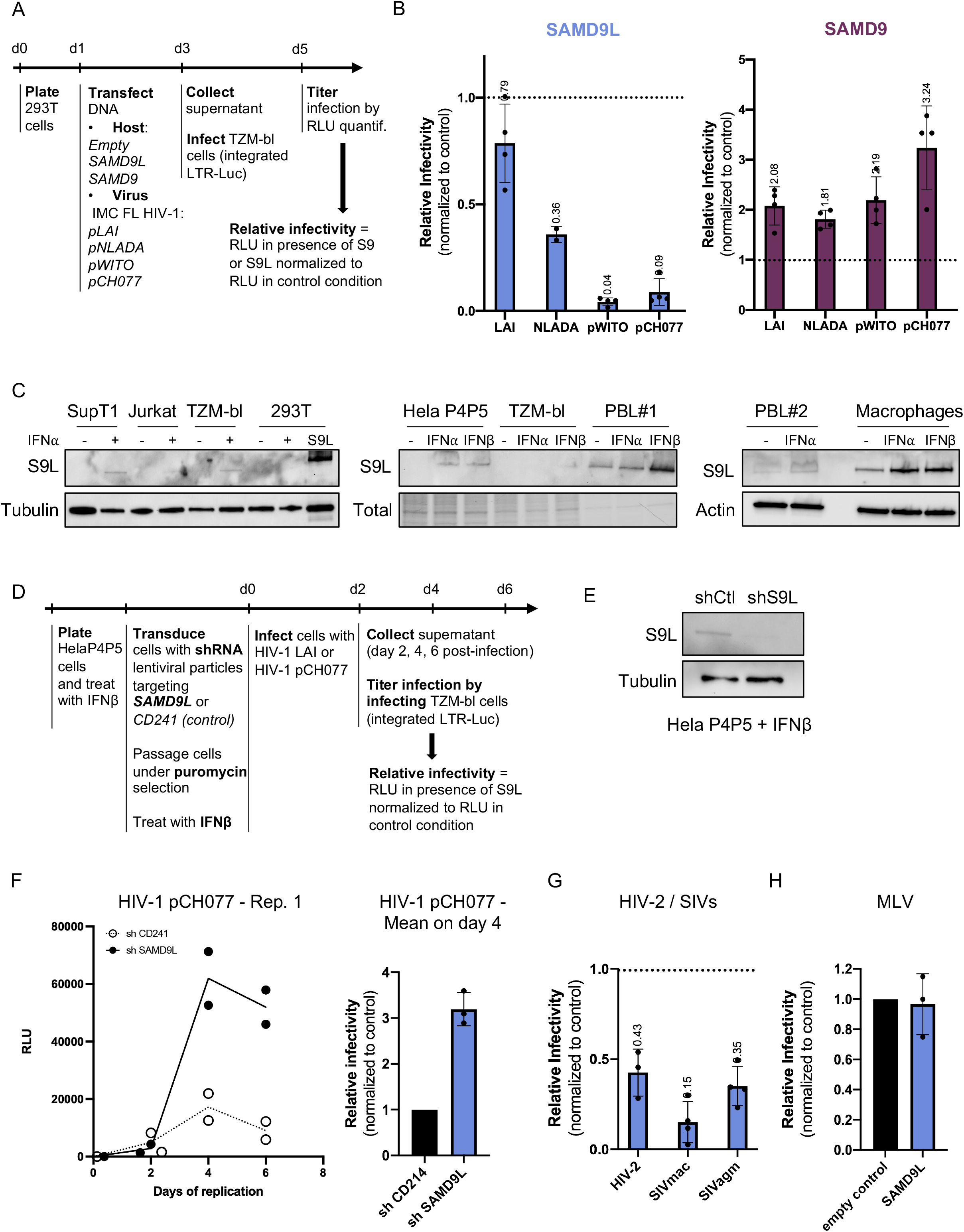
Ectopically-expressed and endogenous interferon-inducible SAMD9L strongly restrict the replication of HIV-1 Transmitted/Founder strains and lentiviruses. A, Experimental setup for the ectopically-expressed SAMD9/9L experiments. IMC FL HIV-1, Infectious molecular clone full-length HIV-1; RLU, Relative light unit; S9, SAMD9; S9L, SAMD9L. Host and virus DNA, and cells as described in Methods. B, Relative infectivity (or infectious yield) of four HIV-1 strains in the context of SAMD9L or SAMD9 expression, normalized to the control in the absence of SAMD9/9L (empty plasmid with the corresponding viral strain). Infectivity was measured as shown in panel A. C, Western-blot analyses of endogenous SAMD9L expression under basal, IFN⍺ or IFNβ stimulation in various cells: immortalized T cell lines (SupT1, jurkat), immortalized fibroblastic cell lines (TZM-bl, 293T, Hela P4P5), primary blood lymphocytes (PBL) from two donors, primary macrophages. The last line of the first blot shows the condition of 293T cells transfected with 1500 ng of TFP-SAMD9L. Loading controls are: Tubulin, total protein (Stain-Free™ gel, BioRad) or Actin. D, Experimental setup for the KD experiments. E, Western-blot of endogenous SAMD9L in HelaP4P5 cells stimulated with IFNβ and transduced with either shCD214 (chCtl) or shSAMD9L. The analysis was performed at day 0 of the viral replication experiment. F, HIV-1 T/F pCH077 replication assay in shCD241 (shCtl) or shSAMD9L HeLaP4P5+IFNβ over 6 days of replication. Titration was performed using the TZM-bl assay. Y-axis represents the raw RLU results for one experiment with the technical duplicates for each condition. Three independent biological replicates were performed for HIV-1 pCH077, pCH077:VSVg, and pLAI (Fig. S3). The mean of three independent biological replicates is shown on the right. G, Same as in 1A-B with HIV-2, SIVmac, and SIVagm IMCs in the context of SAMD9L overexpression (normalized to the control with an empty plasmid). H, Infectivity of retroviral MLV:GFP particles produced with or without SAMD9L. Infectivity was measured by the % of GFP positive cells in 293T target cells (normalized to the control with an empty plasmid).

We next determined, by Western-blot analyses, SAMD9L expression in basal condition and upon type I interferon (IFNα and IFNβ) stimulation, in our experimental settings and in natural HIV target cells. We found that endogenous SAMD9L expression was undetectable in all immortalized cell lines at basal condition (Fig. 1C). Upon IFN stimulation, SAMD9L expression was slightly detectable in SupT1, Jurkat and TZM-bl, and it was enhanced in HelaP4P5 cells (a Hela cell line engineered to express CD4^hi^CCR5^hi^CXCR4^hi^ to serve as HIV target cells) (Fig. 1C). Furthermore, endogenous SAMD9L was expressed in primary macrophages and in primary blood lymphocytes (PBL), albeit with highly varying levels depending on the blood donors (Fig. 1C). Its expression was also enhanced under IFN⍺ or IFNβ stimulation. These results confirm that SAMD9L is an interferon-stimulated gene (ISG)^7^ with limited expression at basal condition.

To determine the functional relevance of endogenous SAMD9L on HIV-1 replication, we silenced SAMD9L expression, by shRNA gene transduction (shSAMD9L and shCD241 as a control ^20^) in HIV-1-permissive cells under IFN stimulation. Because PBL donors exhibited variable SAMD9L expression levels, we used HelaP4P5 cells under IFNβ stimulation for these knock-down (KD) experiments (Fig. 1D-E). KD cells were then infected in technical duplicates with viral titers equivalent to 10ng of p24 Gag of infectious HIV-1 T/F pCH077, HIV-1 LAI, or HIV-1 T/F pCH077:VSVg (i.e. the VSVg-dependent entry is only involved in the first round of replication). We monitored viral replication over a six-day period, with supernatant being collected every two days and titered using TZM-bl reporter cells, and the complete experiment was performed in two to three independent biological replicates (Fig. S3). We found that HIV-1 infectious titers, over a six-day replication curve, were higher in SAMD9L KD cells compared to the KD controls (Fig. 1F, Fig. S3), showing that endogenous IFN-stimulated SAMD9L is an antiviral factor against HIV-1.

Finally, we tested SAMD9L ability to impact the infectivity of several human and simian lentiviruses (simian immunodeficiency viruses, SIVs) beyond the pandemic HIV-1. We found that SAMD9L could also restrict the distantly related HIV type 2, HIV-2 (Fig. 1G). Furthermore, SAMD9L also strongly restricted the infectious yield of replication-competent simian lentiviruses, SIVagm.Tan1 from African green monkey tantalus and SIVmac from rhesus macaque (Fig. 1G), showing that SAMD9L is an antiviral factor of primate lentiviruses.

To test whether SAMD9L broadly affected all viruses, we tested its activity against a gammaretrovirus MLV and two distant RNA viruses: Mopeia virus (MOPV) from the *Arenaviridae* family and Vesicular stomatitis virus (VSV) from the *Rhabdoviridae* family. In the context of SAMD9L exogenous overexpression, we found no significant changes in viral titers during replication (Fig. 1H, Fig. S4; for VSV, we used the antiviral ISG20 as positive control ^21^). This showed a lentivirus-specific antiviral effect of SAMD9L, regarding the other retroviruses and RNA viruses tested in our conditions.

### SAMD9L restricts HIV-1 translation

To identify the lentiviral replication step affected by SAMD9L, we analyzed the levels of proteins secreted in the supernatant (by quantification of Gag p24 or p27 ELISA, and by the quantification of the reverse transcriptase (RT) activity), and we assessed for viral and host proteins in the producer cells (by Western-blot analyses of HIV-1 structural Env and Gag proteins and of SAMD9L) (Fig. 2A). We further kept the SAMD9 condition, as a control of the SAMD9L-specific effect in our experimental settings. Under these conditions, SAMD9L, but not SAMD9, decreased the extent of virion particles released in the cell supernatant (i.e. lower Gag expression and RT activity), which was paralleled by a concomitant decrease in the extent of their intracellular expression (Fig. 2A-C, Fig. S5-6). This effect was even more pronounced for SAMD9L strongly-restricted strains, such as HIV-1 T/F pWITO and pCH077 (Fig. 2B-C, Fig. S5-6). Given that when normalized for released viral protein content, the infectivity of virion particles was not affected by SAMD9L, these results indicate that the major effect of SAMD9L occurred at the level of viral protein accumulation in virion producer cells. To determine whether SAMD9L acted on transcription from viral LTRs or on the newly synthesized HIV-1 RNAs, we performed RT-qPCRs targeting two HIV-1 regions, gag and LTR U5, of the HIV-1 pWITO transcripts in the producer cells. We found that SAMD9L did not modulate HIV-1 RNAs, nor cellular U6 and TBP mRNAs used as controls (Fig. 2D), showing no defect on viral or cellular RNAs and implying a post-transcriptional effect.

**Figure 2.**
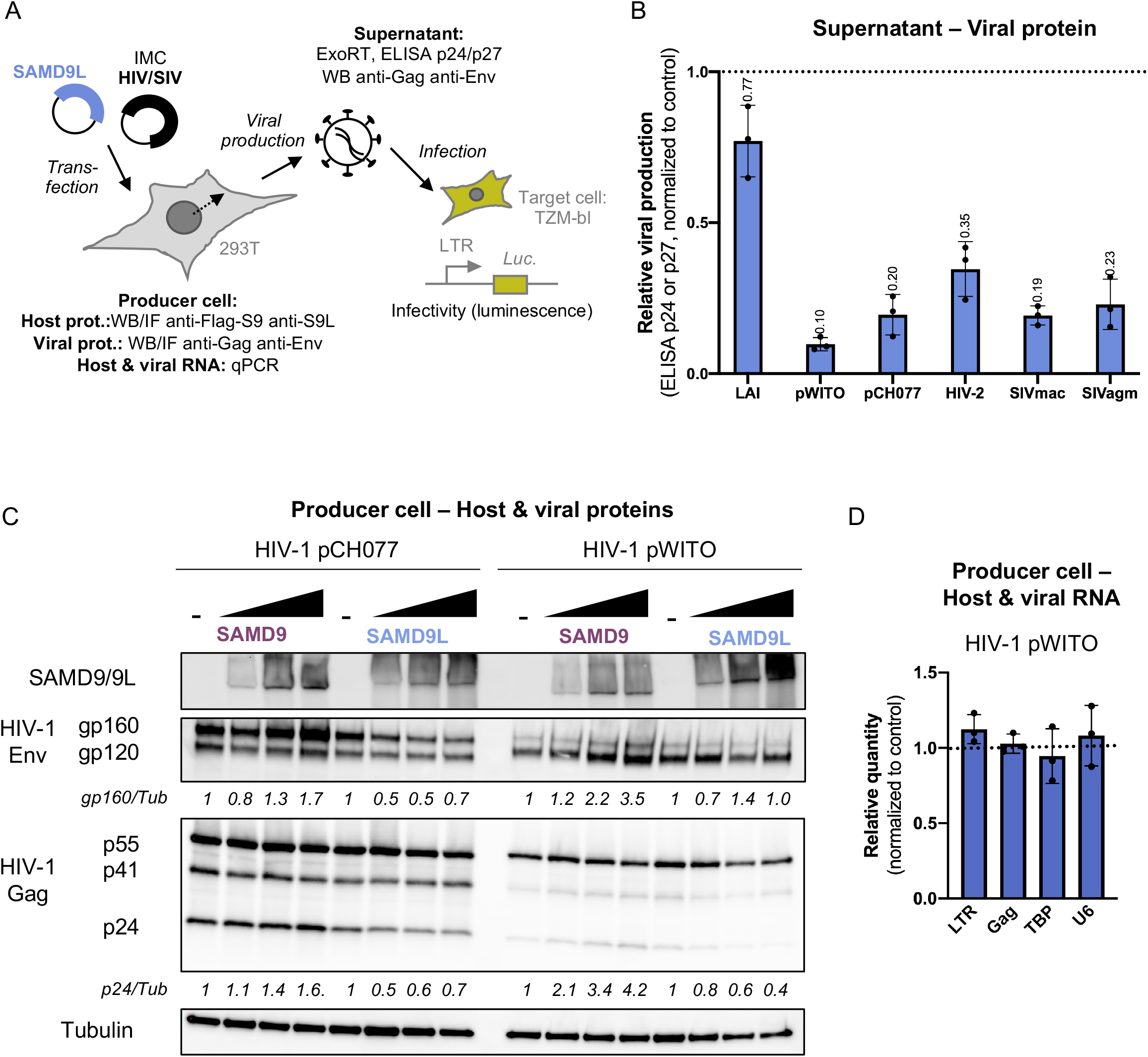
SAMD9L restricts HIV and lentiviruses at the translation step. A, Experimental setup. Set up similar to Fig. 1A with detailed analyses to investigate the viral restriction. B, Relative viral production of diverse primate lentiviruses, as measured by ELISA p24 (for HIV-1 strains) or p27 (for HIV-2 and SIVs) in the supernatant of producer cells, normalized to the control in the absence of SAMD9L. C, Western-blot analyses of exogenously-expressed SAMD9 (anti-FLAG) or SAMD9L (anti-SAMD9L) proteins, as well as of lentiviral protein Env and Gag in HIV-1 producer 293T cells. Below the lanes, quantifications are provided for gp160 and p24 protein expressions normalized to Tubulin, and expressed as fold difference compared to the condition in the absence of SAMD9/9L (normalized to 1). Western-blot experiments were performed in multiple replicates and for all the HIV-1, HIV-2 and SIV strains (Fig. S5-6). D, Relative RNA quantity in the HIV-1 producer cell for HIV-1 LTR and Gag targets, as well as cellular TBP and U6, in the presence or absence of SAMD9L. Results are normalized to the control in the absence of SAMD9L.

Lastly, we tested whether SAMD9L could also impact the early phases of HIV-1 and SIVmac replication, by using single-round non-replication-competent viruses (HIV-1 and SIVmac viral-like particles pseudotyped with VSVg and encoding for a firefly luciferase reporter gene in the transfer plasmid, HIV-1:Luc and SIVmac:Luc, respectively) in infections of target cells overexpressing SAMD9L. We found that target cells overexpressing SAMD9L had similar levels of lentiviral infection compared to control cells (Fig. S7), showing that SAMD9L does not impact the early phases of lentiviral replication.

Therefore, we show that SAMD9L restricts HIV-1 and other lentiviruses’ replication by affecting viral translation, with a stronger effect against HIV-1 T/F strains compared to the HIV-1 laboratory-adapted strains.

### HIV and lentivirus restriction by SAMD9L depends on E198/D243 in a SLFN-like active site

SAMD9 and SAMD9L are large proteins predicted to have a similar and complex domain architecture (Fig. 3A)^22^. The recently solved crystal structure of the SAMD9 156-385 amino acid region showed it can bind DNA^12^. This region corresponds to the AlbA2 domain, which typically binds DNA and/or RNA and possesses multimerization properties^22,23^. Before the SAMD9(156-385) structure was available (Fig. S8A), we had analyzed SAMD9L with the protein homology and structure prediction tool HHpred and found a homology with members of the Schlafen (SLFN) gene family (Fig. S8B). This homology localized into their respective AlbA2, which is present in all SLFNs and forms the C-lobe of their N-terminal domain, also known as the SLFN-box ^24^ (Fig. 3A). SLFN genes are involved in multiple cellular processes, including development, cancer and antiviral immunity ^24^. The SLFN-box presents nucleic acid binding properties, as well as endoribonuclease activity in several SLFNs, and the enzymatic active site is dependent on three conserved acidic amino acids (two Glu (E) and one Asp (D)) ^24,25^. Interestingly, SLFN11 and SLFN13 restrict HIV-1 replication by inhibiting translation and this enzymatic active site is necessary for this function ^25–28^. We therefore compared the structures of the SAMD9-AlbA2 domain, the AlphaFold-predicted SAMD9L, and the SLFN-box of SLFN5, SLFN12 and rSLFN13. We found strong structural homology and conservation of the active site triad in SAMD9/9L: E184/E196/D241 in SAMD9 and E186/E198/D243 in SAMD9L (Fig. 3B-C, Fig. S8). A protein sequence alignment of SLFNs with SAMD9/9Ls further showed a conservation of these three residues in SAMD9/9L mammalian sequences (Fig. 3D). We therefore suspected their key role in SAMD9L antiviral function against lentiviruses.

**Figure 3.**
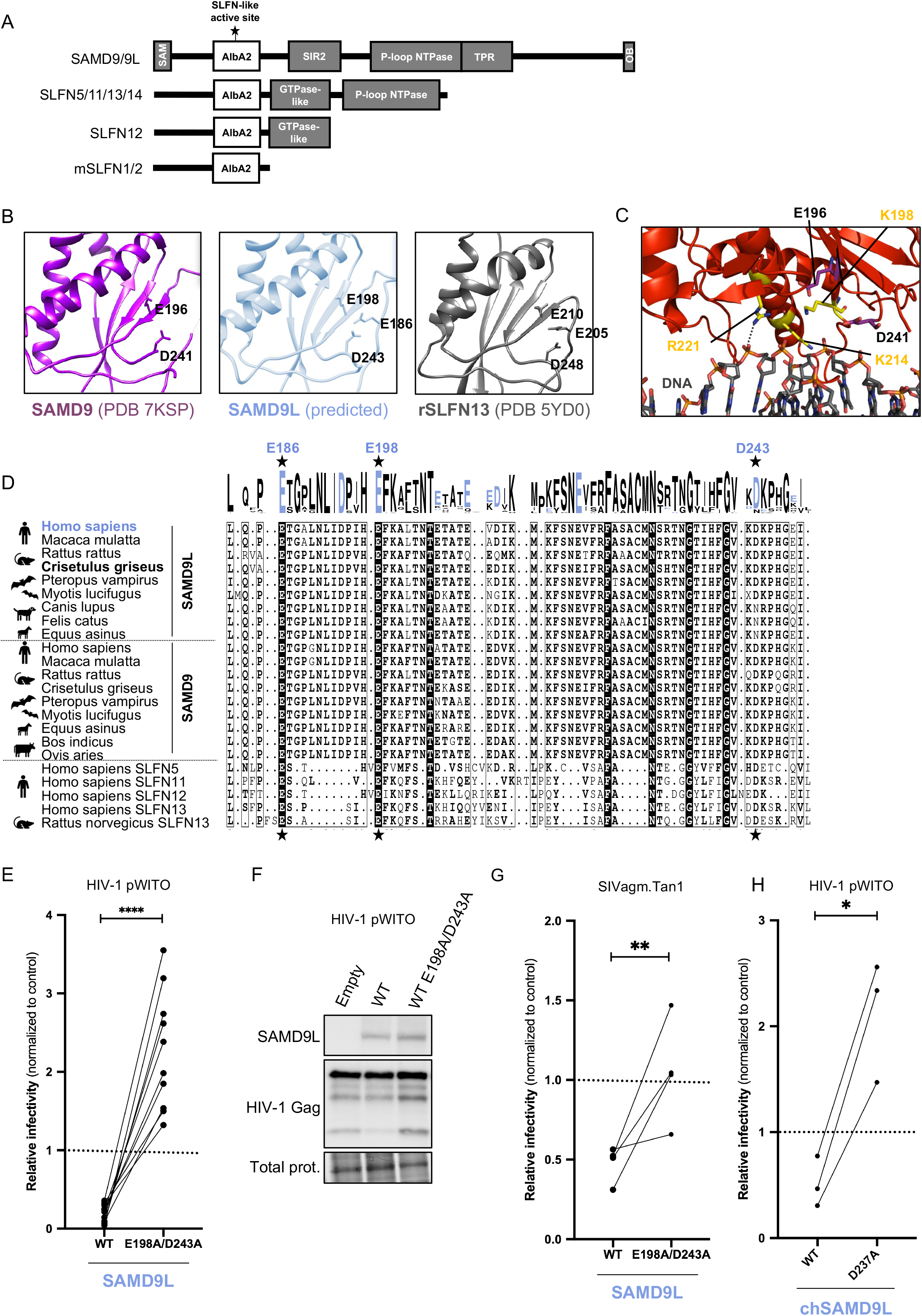
SAMD9L restriction of lentiviral translation is governed by residues E198/D243 in a Schlafen-like box active site, conserved in mammals. A, Representation of the protein domains in SAMD9/9L and SLFNs with the identification of the putative active site in a SLFN-like box. mSLFN1/2, mouse SLFNs. B, Structure of the SLFN-like box in SAMD9, SAMD9L and rSLFN13 (rat SLFN13). Atoms of the residues forming the active site are represented with sticks and their coordinates labeled in black. SAMD9 and rSLFN13 are from solved crystal structures (PDB 7KSP and 5YD0, respectively), while SAMD9L is from AlphaFold prediction. C, SAMD9 crystal structure in complex with DNA (PDB 7KSP). Atoms of the residues forming the active site are represented with purple sticks (labels in black), while those involved in DNA binding are in yellow. D, Amino-acid sequence alignment of the region bearing the SLFN-like box in SAMD9/9L and SLFNs from various mammalian species. The triad of residues forming the active site is labeled with black stars with coordinates from human SAMD9L above in blue. In the alignment: Residues with black font are strictly conserved among the sequences; Dots correspond to alignment gaps; Bold residues are the one matching with the consensus. The logo plot is from WebLogo with the acidic residues (Asp, D and Glu, E) in blue. E, Relative infectivity of HIV-1 pWITO in empty, SAMD9L or SAMD9L-E198A/D243A conditions, following Fig 1A experimental design. F, Western-blot analysis in the corresponding producer cells showing SAMD9L WT and mutant expression, as well as HIV-1 Gag proteins. Loading control is from total protein (BioRad Stain-Free gel). G, Similar to panel E with SIVagm.Tan1. H, Similar with the effect of Chinese hamster chSAMD9L WT and D237A on HIV-1 pWITO infectivity. Statistics: ****, p value < 0.0001. **, p value < 0.01. *, p value < 0.05.

To test this, we mutated the active site triad at two residues generating the SAMD9L-E198A/D243A mutant plasmid, and we assessed its effect on HIV-1 pWITO and SIVagm.Tan1. We found that the HIV-restriction ability of SAMD9L was totally abolished by the E198A/D243A mutations (Fig. 3E). In accordance, the expression of HIV-1 structural proteins in the cell was higher in the presence of SAMD9L-E198A/D243A compared to wild-type (WT) SAMD9L (Fig. 3F). Furthermore, SAMD9L antiviral effect against SIVagm.Tan1 was also dependent on the E198/D243 site (Fig. 3G).

To determine if the antiviral function and the motif were also conserved and important in another mammalian SAMD9L ortholog, we tested the Chinese hamster SAMD9L ^29^ and its respective mutant in the predicted active site, chSAMD9L-D237A. We found that chSAMD9L also restricted HIV-1 pWITO and that chSAMD9L-D237A lost its anti-lentiviral capacity (Fig. 3H), showing that this catalytic site is also necessary in Chinese hamster SAMD9L for lentiviral restriction.

Altogether, we identified and showed that E198 and D243 from the active site of the SLFN-like box are necessary for SAMD9L anti-lentiviral restriction.

### Mutating the SLFN-like active site, E198/D243, also relieves the pathogenic translational inhibition of SAMD9L-F886Lfs*11 from SAAD/ATXPC patients

Germline gain-of-function (G-o-F) mutations in SAMD9L are responsible for diverse multisystem disorders, including myeloid malignancies and systemic autoinflammatory diseases in humans^2,6–8^. These mutations are predominantly localized in its C-terminal half-part, mostly within the putative P-loop NTPase domain^8^. One such variant from SAAD/ATXPC patients is SAMD9L-F886Lfs*11, a germline frameshift mutant that generates a C-terminal truncated protein^2^ (Fig. 4A). It is associated with a G-o-F phenotype, dramatically suppressing cellular protein synthesis^1–3^. The basic residues (K or R) in the SAMD9L AlbA2, which are involved in dsNA binding, are necessary for this translation inhibition^12^. To determine the importance of the SLFN-like acidic E198/D243 residues (Fig. 4A) on cellular translation inhibition, we performed the Click-iT™ Plus O-propargyl-puromycin (OPP) Synthesis Assay, by overexpressing in 293T cells SAMD9L WT, SAMD9L-F886Lfs*11 (i..e. SAMD9L truncation based on the patient’s natural variant), or their corresponding mutants E198A/D243A and by measuring OPP incorporation in nascent cellular proteins by flow cytometry. We showed that the mild translational repression by SAMD9L WT and the strong inhibition exerted by the SAMD9L-F886Lfs*11 were strictly dependent on E198/D243 (Fig. 4B-C). Therefore, E198/D243 of the SLFN-like box active site in SAMD9L are essential and drive cellular protein synthesis repression.

**Figure 4.**
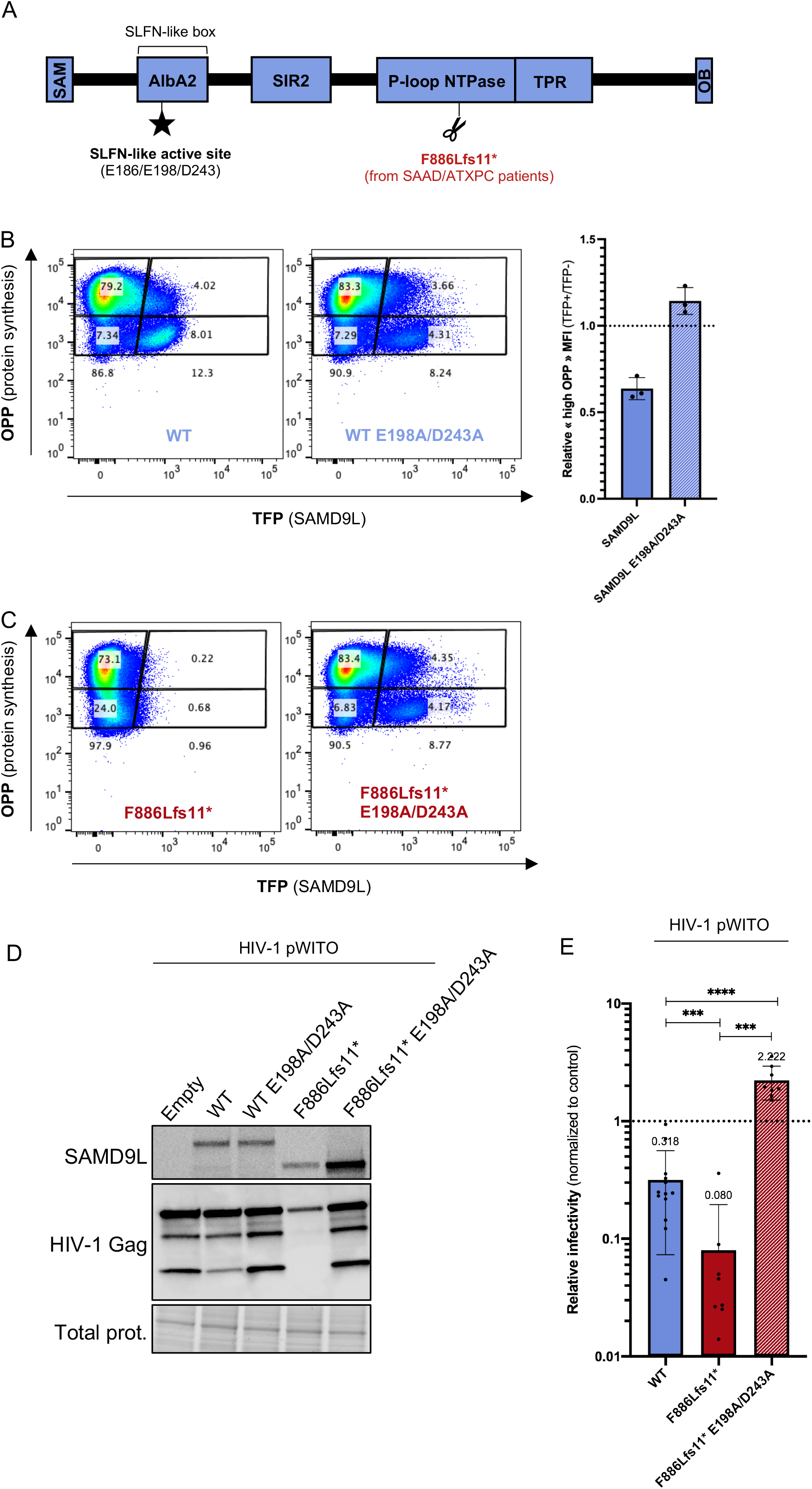
The cellular translational repression of SAMD9L and SAAD/ATXPC patient variants is alleviated by the E198A/D243A mutations in the SLFN-like box. A, Representation of the protein domains in SAMD9L highlighting the truncation of one patient variant, SAMD9L-F886Lfs11* ^2^, and the SLFN-like box active site identified in this study. B, Representative flow cytometry density plots of OPP incorporation relative to the expression of either TFP-SAMD9L WT or WT E198A/D243A. Plots are divided into four polygons representing TFP+ cells (right), TFP-cells (left), high OPP cells (up) and low OPP (down). Corresponding cell percentages are labeled into corresponding polygons and sums of TFP+ and TFP-cells are labeled below. On right, quantification from three independent biological replicates of the protein synthesis assay. For each replicate, the relative “high OPP” MFI was calculated using the MFI of the “high OPP” TFP+ polygon normalized to the MFI of the “high OPP” TFP-polygon. MFI, median fluorescence intensity. C, Similarly, effect of F886Lfs11* and F886Lfs11* E198A/D243A. Of note and as previously reported, SAMD9L-F886Lfs11* also represses its own protein synthesis. D-E, Effect of SAMD9L patient variants and E198/D243 mutants on HIV-1 protein expression (D) and relative infectivity (E). Experimental conditions are as shown in Fig. 2A. ****, p value < 0.0001. ***, p value < 0.001.

Furthermore, we tested how SAMD9L-F886Lfs*11 impacted lentiviral replication and found that it strongly inhibited HIV-1 protein expression and infectivity, increasing the antiviral effect by five-fold compared to SAMD9L WT (Fig. 4D-E). This viral restriction was also fully abolished by the E198A/D243A mutations. Therefore, the pathogenic G-o-F SAMD9L-F886Lfs*11 exerts an increased anti-lentiviral activity, and importantly, the exacerbated shutdown of both cellular and lentiviral protein synthesis can be relieved by mutating the SLFN-like active site.

## Discussion

In this study, we identified SAMD9L, but not SAMD9, as an IFN-inducible anti-lentiviral factor that is most potent against HIV-1 Transmitted/Founder patient strains, but does not affect the gammaretrovirus MLV or distant RNA viruses VSV and MOPV. Mechanistically, we showed that SAMD9L restricts lentiviruses at the translation step. Furthermore, we discovered SAMD9L determinants of lentiviral and cellular translational repression: an essential SLFN-like box effector domain with key E198/D243 residues, and a C-terminal auto-regulatory domain, previously identified in the context of SAAD/ATXPC patients. Altogether, our study links antiviral and cellular properties and enlightens key effector determinants of SAMD9L variants involved in severe autoinflammatory or ATXPC diseases.

We describe the identification of human SAMD9L as one effector of the anti-HIV activity of type I IFN response, in the late phases of replication. It may have been missed from most previous HIV-host factor screens, because SAMD9L is poorly expressed in most immortalized cell lines used in these screens (except THP-1 myeloid cells in which Oh Ainle et al identified SAMD9L as a potential candidate^17^), and because SAMD9L antiviral effect is the strongest against HIV-1 T/F strains, which are rarely tested in first approaches. Understanding the molecular details for SAMD9L auto-regulation, the viral determinants involved in the lentiviral strain-specificity, and the basis for the dichotomy between SAMD9L/HIV-1 inhibition and SAMD9/HIV-1 enhancement will help to elucidate the mechanism of this cell-autonomous immune defense function. As HIV-1 T/F strains are more susceptible than laboratory adapted strains, it would also be interesting to perform within-patient longitudinal analyses of HIV-1 strains during infection regarding their SAMD9L susceptibility to determine whether HIV-1 evolves and overcomes SAMD9L restriction over time and by which mechanism. SAMD9 and SAMD9L have both evolved under positive selection in mammals^30^, reflecting potential adaptation to past viral epidemics, which is a typical feature of host restriction factors and viral interacting proteins^31–33^. These past “evolutionary arms-races” may have involved conflict with ancient poxviruses^9^, as well as lentiviruses and possibly other viral families.

We identified key acidic residues E186/E198/D243 in the AlbA2 domain of SAMD9L, which are similar to the SLFN box and could constitute a ribonuclease active site. The recent crystal structure of SAMD9 AlbA2 solved in complex with DNA^12^ showed the capacity of SAMD9/9L to bind dsNA. One primary hypothesis is therefore that SAMD9L AlbA2 effector domain may bind and cleave tRNAs through the E186/E198/D243 active site, similarly as SLFN13 or SLFN11^25,27,34^. Interestingly, SAMD9L and SLFN11 are also both involved in DNA damage response, a function mediated by type II tRNAs cleavage in SLFN11^1,34^. Other hypotheses are that SAMD9L effect may be linked to mRNA stability or ribosomal protein disruption^35^, or given the electrostatic environment of E198/D243, it is possible that the residues convey a proper orientation for R221 allowing its interaction with the dsNA phosphate backbone for efficient binding^12,25^. This latter would be more similar to SLFN5, which only binds tRNA without cleavage^36^. Related to these mechanisms, an intriguing possibility is that SAMD9L could act in a codon usage dependent manner, inhibiting viral and host genes with strongly biased codon usage (such as poxviruses or lentiviruses, but not MLV retrovirus)^37–40^, which could drive SAMD9L viral-specificity and the deleterious effect of SAMD9L G-o-F variants. A codon usage similarity analysis of virus transcripts that are resistant or susceptible to SAMD9L restriction (Fig. S9) shows a strong correlation with SAMD9L antiviral-specificity (Fig. 5), suggesting that codon bias may, at least partly, underlie SAMD9L translational repression mechanism.

**Figure 5.**
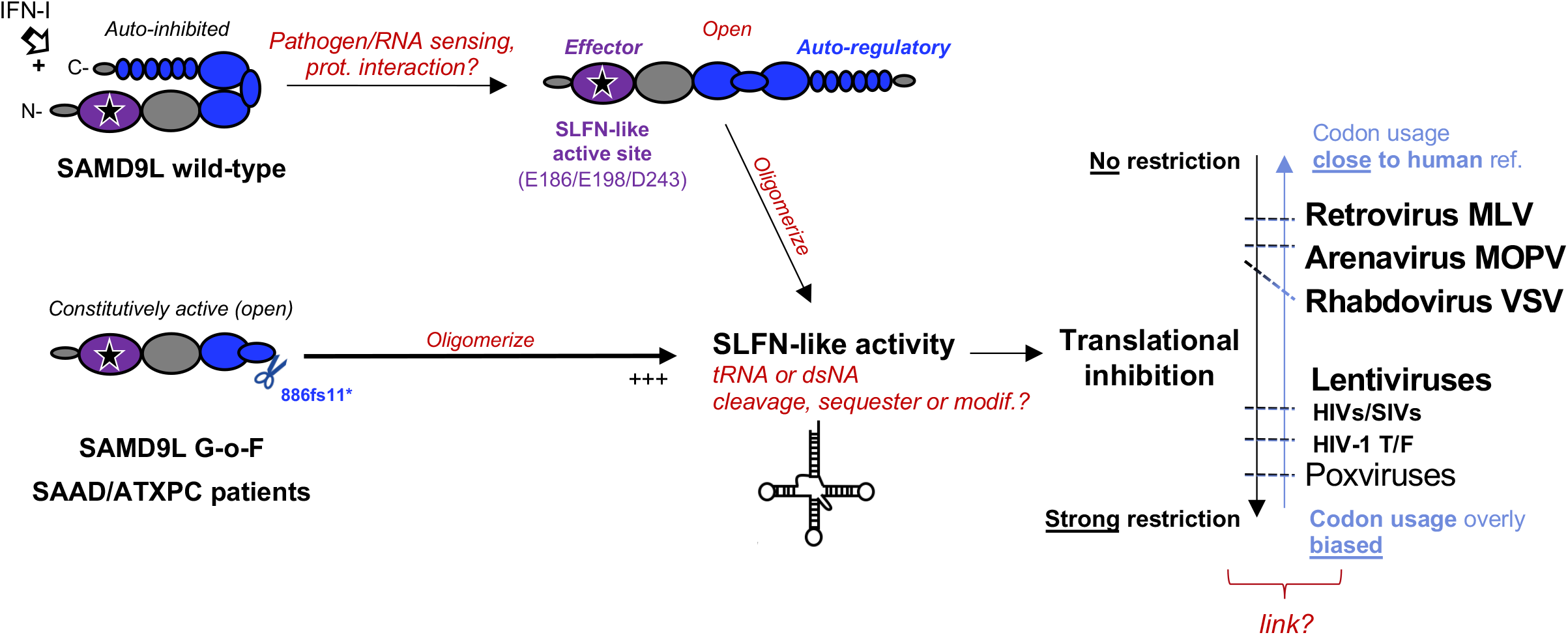
Key findings and proposed model of SAMD9L antiviral and cellular functions. Model for SAMD9L in normal condition (wild-type) and in SAMD9L G-o-F variant F886fs11* from SAAD/ATXPC patients. The AlbA2 effector domain is in purple with the black star representing the SLFN-box active site. The P-loop NTPase and TPR auto-regulatory domains are in dark blue (with a tripartite representation of the P-loop NTPase and with TPR represented by repeated adjoining circles). Texts in red correspond to hypotheses. The antiviral restriction metric is from this study (Retrovirus, Lentivirus, Arenavirus, Rhabdovirus) and previous findings (Poxvirus). The codon usage metric corresponds to Fig. S10. We interrogate a potential link between (i) strong translational inhibition of viral and cellular transcripts and (ii) codon usage bias (overly different to the human reference).

Whatever its exact function, the identification of these host determinants in SAMD9L is of additional primary importance, as we show that targeting them allows to revert the effect of the pathogenic G-o-F SAMD9L variants from patients with SAAD or associated myeloid disorders, which would have direct clinical relevance.

SAMD9 and SAMD9L are both antiviral against poxviruses, whereas only SAMD9L is antiviral against HIV. While sharing only 60% amino acid similarity, the paralogs harbor high structural homology. A hypothesis to explain their divergent effect regarding HIV may reside in the escape of SAMD9 antiviral activity through an HIV antagonist or through HIV evasion. Because we observe a slight proviral effect of SAMD9 on HIV-1, it would be interesting to investigate whether HIV-1 has even adapted to usurp the IFN-inducible SAMD9 as a host co-factor. This may be similar to influenza virus, which is able to repurpose the IFN-induced IFIT2 to promote viral protein translation^41^. Alternatively, there may be intrinsic functional differences between SAMD9 and SAMD9L, for example it is possible that SAMD9L is able to sense and/or restrict HIV, as opposed to its paralog. In these cases, during the SAMD9/9L divergence, paralogous genes may have (i) retained similar functions against poxviruses, allowing increased antiviral potency – such as observed for the antiviral IFITM2 and IFITM3^42,43^ or for the anti-retroviral primate APOBEC3s^44,45^, and (ii) specialized, with SAMD9L bearing anti-lentiviral functions, allowing antiviral breadth and diversification of the effector functions – as observed for the mammalian MX, IFIT, or SLFN gene families^13,24,43,46,47^.

Further structural homology analyses on full-length SAMD9L allowed us to identify, in the intermediate/C-terminal region, a homology with the protein PH0952 from an archaea, *Pyrococcus horikoshii*, harboring a double-inhibition mechanism with (i) a signal transduction ATPase with numerous domains (STAND), analogous to Apoptotic peptidase activating factor 1 (APAF-1) as previously reported^3,22^ and (ii) a tetratricopeptide repeat (TPR) sensor domain^48^. Thus, the intermediate region of SAMD9L would include at least two regulatory mechanisms, possibly self-guarding an accidental activation or exposure of the SLFN-like box (Fig. 5). These types of regulatory mechanisms may have ancient evolutionary roots shared between eucaryotes, archaea and bacteria^49^. In accordance with previous models, the F886LFs11* truncation would lead to a constitutive cellular^3^ and antiviral translational shutdown. Therefore, while the N-terminal region contains the necessary and sufficient effector domain for SAMD9L-driven translational repression, the C-terminal region could provide a strong self-guarding, whose mechanism and evolutionary origin remain to be characterized.

Although distantly related, interferon-stimulated SAMD9/9L and SLFN proteins bear some redundant and crucial cellular and innate immune antiviral functions. Both proteins are differentially, but tightly, auto-regulated like many genes involved in Type I interferonopathies^16^. Malfunction in these gene families can lead to dramatic human genetic and infectious diseases and underlie a determinant role for their proper regulation in the cell, which require further mechanistic and translational research.

## Materials and Methods

### Cell lines and culture

Human embryonic kidney 293T (ATCC, cat. CRL-3216), HelaP4P5 (Hela cells expressing CD4^hi^CCR5^hi^CXCR4^hi^), and TZM-bl (NIH AIDS Research and Reference Reagent Program, Cat. 8129) cells were maintained in Dulbecco Modified Eagle Medium (DMEM) supplemented with 10% fetal calf serum (FCS, Sigma cat. F7524) and 100 U/ml of penicillin/streptomycin. TZM-bl cells express CD4, CCR5 and CXCR4, and encode for Luciferase and β-galactosidase under the LTR promoter. They are routinely used for titration of lentiviral supernatants. SupT1 and Jurkat T cells were maintained in RPMI-1640 with 10% FCS. Primary blood monocytes and lymphocytes were purified from leukopacks of healthy blood donors, using Ficoll and Percoll gradients. Lymphocytes were activated for >24h with 1µg/ml PHA (Sigma) and 150 U/ml IL2 (Eurobio, PCUT-209). Monocytes were purified using Miltenyi monocyte isolation kit II (cat. 130-091-153) and differentiated in macrophages upon incubation for >4 days in complete RPMI-1640 with 10% FCS and 100 ng/ml Macrophage-Colony Stimulating Factor (M-CSF, Eurobio, cat. 01-A0220-0050).

### Plasmids

The virus and host plasmids used in this study are shown in Table S1. SAMD9L-E198A/D243A, SAMD9L-F886Lfs*11, SAMD9L-F886Lfs*11-E198A/D243A were generated from pMAX:mTFP1-SAMD9L plasmid using the QuikChange Lightning Site-Directed Mutagenesis Kit (Agilent) following the manufacturer’s instructions. Similarly, chSAMD9L-D237A was generated from the original chSAMD9L plasmid (pcDNA3.1/V5-His-Topo:3xFLAG-chSAMD9L). The frameshift F886Lfs*11 was generated by deleting two threonines.

### Viral-like particle (VLP) production and infection for single-round GFP-or Luciferase-reporter HIV-1, SIVmac, and MLV. Production

293T cells were transfected by calcium phosphate method with DNA plasmids encoding the packaging plasmid, the genome plasmid, and the envelope plasmid (VSVg, pMD2.G) at ratios of 2:2:1. For HIV-1 VLPs, the packaging plasmid was pHIV-1 GagPol 8.2 (that includes most accessory genes) or psPAX2 (devoided of accessory genes), and the reporter genome plasmid was pNaldini-GFP or pHIV-1-LTR-fLuc. For SIVmac VLPs, the packaging plasmid was pSIV3+ (that includes most accessory genes), and the reporter genome plasmid was pGAE-fLuc. For MLV VLPs, the packaging pMLV plasmid was pTG5349 and the reporter plasmid pMLV-CMV-GFP was pTG13077. Details and references of plasmids are available in Table S1. Two to three days post-transfection, the viral supernatant was filtered through a 0.45 ⌈M diameter filter or purified by ultracentrifugation. Supernatant was then used for titration by exogenous RT activity (10 µl), or p24/p27 ELISA (10 µl), or TZM-bl assay (25 µl), and used for subsequent infection.

### Infection

Different amounts of purified VLPs were used to infect 293T cells, which were transfected 24h or 48h before with either empty, SAMD9 or SAMD9L plasmid. For GFP-VLPs, cells were trypsinized two days post-infection and fixed in PFA before quantification on BD FACSCanto II (SFR BioSciences) and analyses in FlowJo. For Luc-VLPs, cells were lysed with BrightGlow Lysis Reagent (Promega E2620) and the mean relative luminescence units (RLU) were directly measured by FLUOStar Optima reader.

### Replication-competent lentivirus production and infection

293T cells were seeded in 6-well plates at 0.2M cells/ml. Twenty-four hours later, cells were co-transfected with TransIT-LT1 (Mirus) with a plasmid encoding a fully replication-competent lentivirus (IMCs), as well as a plasmid encoding either SAMD9, SAMD9L or an empty control. Unless stated in the text or figure (e.g. for viral and host doses), the DNA quantity was 1500 ng for the host plasmids and 1200 or 1600 ng for the virus plasmids. Forty-eight hours post-transfection, the cells were harvested for western-blot or qPCR analyses. The supernatants were collected and stored at -80ºC for further analyses, including titration by ELISA p24/p27 or by RT assay, Western-blot analysis after ultracentrifugation, or infection of TZM-bl cells. For infection, TZM-bl cells were seeded in 96-well plates and infected by a serial dilution of viral supernatant. Forty-eight hours post-infection, cells were lysed using BrightGlow Lysis Reagent (Promega E2620) and the RLU were measured by Tecan Spark® Luminometer. Infectivity of viruses in various conditions are always expressed as fold-change as compared to a paired viral infection condition in the absence of any SAMD9/9L.

### Western blotting

Cells were harvested, lysed by ice-cold RIPA buffer (50 mM Tris pH8, 150 mM NaCl, 2 mM EDTA, 0.5% NP40) supplemented with protease inhibitors (Roche) and by sonication. For the supernatant fraction, 1 ml of HIV-1 replication-competent supernatant was collected and purified through sucrose by ultracentrifugation, and the pellets were resuspended in ice-cold RIPA buffer. Proteins from cell lysates or from supernatant were then similarly processed: separated by electrophoresis and then transfected on PVDF membrane through a wet transfer overnight at 4°C. The membranes were blocked in a TBS-T 1X solution (« Tris Buffer Saline », Tris HCl 50 mM pH8, NaCl 30 mM, 0.05% of Tween 20) with 5% powder milk and were incubated for 1h to overnight in primary antibodies and 1h in secondary antibodies. Detection was made with SuperSignal West Pico Chemiluminescent Substrate (ThermoFisher Scientific) usinq the Chemidoc Imagina System (Biorad). For endogenous expression of SAMD9 and SAMD9L, cells were stimulated, or not, with IFNα2 (Tebu Bio, cat. 11100-1) or IFNβ (500 IU /ml, R&D systems, 8499-IF-010) for 24h prior lysis. The following antibodies were used: anti-Tubulin (Sigma, cat. T5168), anti-Actin (Sigma, cat. A2228) anti-Flag (Sigma, cat. F3165), anti-SAMD9L (Proteintech, 25173-1-AP), anti-SAMD9 (Sigma, HPA021319), anti-Gag (NIH HIV Reagent Program, 183-H12-5C), anti-Env (Aalto, D7324), secondary anti-mouse and anti-rabbit IgG-Peroxidase conjugated (Sigma, cat. A9044 and AP188P, respectively). Total protein was also used as loading control, using the PrestainedTM BioRad gels.

### Virus release titrations by ELISA or RT assays

Titration of HIV-1, as well as HIV-2 and SIV, viruses in the supernatant were performed using HIV-1 p24 and SIV p27 ELISA kits (XpressBio), respectively, following the manufacturer’s instructions. Reverse transcriptase RT activity of retroviruses in viral supernatant was determined with an exogenous RT reaction, in which the ability of RT to incorporate ^32^P-labeled dTTP into a poly(rA)/oligo(dT) matrix is quantified ^50^.

### Cell counting

293T cells were seeded in 6-well plates at 0.2M cells/ml and were transfected with TransIT-LT1 (Mirus) 24h later with 1500 ng of empty plasmid, or of plasmid encoding SAMD9 or SAMD9L, as in other experiments. Cells were counted with trypan blue at 0, 24, 48 and 72h post-transfection, corresponding to the maximum period during which the cells are maintained for virus replication experiments. The experiment was performed in triplicates in two independent biological replicates.

### HIV-1 replication in SAMD9L knock-down cells

The shRNA oligos were designed from the Broad Institute Genetic Perturbation platform and the shRNA clones in pLKO.1 were purchased from Sigma. The three oligo sequences for pLKO.1-shSAMD9L are: 5’-CCGGGCAGACAGTATTGCACTAAATCTCGAGATTTAGTGCAATACTGTCTGCTTTTTTG, 5’-CCGGCATCGCTACATAGAACATTATCTCGAGATAATGTTCTATGTAGCGATGTTTTTTG, 5’-CCGGGCTCTTATGTTACTGACTCTACTCGAGTAGAGTCAGTAACATAAGAGCTTTTTTG. shRNA oligos targeting CD241 (also named RHAG) were chosen as non-target controls: 5’-CCGGGCAAGAATAGATGTGAGAAATCTCGAGATTTCTCACATCTATTCTTGCTTTTTG, 5’-CCGGCCTCTGACATTGGAGCATCAACTCGAGTTGATGCTCCAATGTCAGAGGTTTTTG, 5’-CCGGGATGACAGGTTTAATTCTAAACTCGAGTTTAGAATTAAACCTGTCATCTTTTTTG. shRNA-coding lentivectors were prepared in 10-cm dishes by transfection of 293T cells, using calcium phosphate method, of 2.5 µg of psPAX2, 0.75 µg of VSVg, and 1 µg of each shRNA-coding construct. After 72h, the supernatant was collected and purified by ultracentrifugation, and the shRNA lentiviral stock was titrated by HIV-1 p24 ELISA (XpressBio). To generate HelaP4P5 SAMD9L KD and control (CD241) KD, the cells were plated at 0.2M cells/ml in a 6-well plate and stimulated with IFNβ (500 IU/ml, R&D systems cat. 8499-IF-010). At 24h, cells were transduced with 100 ng p24 of shRNA lentiviral particles targeting SAMD9L or CD241. After 48h, cells were then passaged under puromycin selection. The expression of SAMD9L in shSAMD9L and shCD241 cells was controlled by Western-blot before HIV-1 replication. For HIV-1 replication, shSAMD9L and shCD241 cells were seeded in 12-well plates, with IFNβ treatment, and were infected in duplicates the next day with replication-competent HIV-1 LAI or HIV-1 pCH077 or HIV-1 pCH077 pseudotyped with VSVg for the first round at 10 ng p24 (HIV-1 p24 ELISA, XpressBio). Media was changed 6h post-infection. Supernatant was then collected at day 0, 2, 4 and 6 post-infection and titrated by infection of TZM-bl in LTR-Luc reporter assay. Each condition was performed in duplicates in a total of two to three independent biological replicates.

### Protein synthesis assay

293T cells were seeded at 0.3M cells/ml in 12-well poly-L-lysine coated plates. Twenty-four hours later, medium was changed, and cells were transfected with TransIT-LT1 (Mirus) with 3 µg of vector encoding SAMD9L or pcDNA3.1 empty. Seventy-two hours post-transfection, cells were incubated in OPP (Immagina Biotechnology) for 30min at 37°C. Medium was discarded, and cells were trypsinized, harvested and fixed with PFA 4%. Cells were then washed with PBS BSA 3% and permeabilized in PBS 0,5% Triton X-100 for 15min. Click-iT® Plus Alexa Fluor® Picolyl Azide assay was then performed and cells were analyzed on MACSQuant® VYB Cytometer (Miltenyi Biotec, SFR BioSciences).

### RT-qPCR

293T cells prepared for replication-competent lentiviral production and infection were harvested and lysed by TRIzol Reagent (ThermoFisher Scientific). Total RNA was extracted and purified following the manufacturer’s protocol. RNA was treated with RQ1 RNase-Free DNase (Promega) following the manufacturer’s protocol. The RT step was performed using the SuperScript® III First-Strand Synthesis System (Invitrogen). FastStart Universal SYBR Green Master (Roche) was used for qPCR and was performed on StepOnePlus™ Real-Time PCR System (ThermoFisher Scientific, SFR BioSciences). Primer sequences were designed to target Gag and LTR from lentiviruses and TBP and U6 from the host (as controls); sequences are available in Table S2. For Gag and LTR RNA quantification, double normalization was performed using TBP and U6. For TBP quantification, U6 normalization was used.

### Structure homology analyses

HHpred was used for homology detection analyses ^51^. Protein structures were retrieved from AlphaFold DB and RCSB PDB ^52–54^. Structural analyses and visualization were performed using UCSF Chimera (The Regents) and PyMOL Molecular Graphic Systems (Schrödinger, LLC) softwares.

### VSV infection

293T cells were seeded in 12-well plate and were transfected 24h later with 800 ng of SAMD9L-encoding plasmid, ISG20-encoding plasmid ^21^ as a positive control for VSV restriction, or an empty plasmid as a negative control. Twenty-four hours post-transfection, cells were infected with VSV-GFP ^55^ at MOI 0.3 and, 16 hours post-infection, the viral supernatant was collected for each condition and titered on new cells through the GFP reporter. Single-cell analysis was acquired with BD FACSCanto™ II Flow Cytometer (SFR BioSciences). The percentage of VSV-GFP+ cells in the absence of SAMD9L or other host factor (empty control) was used as the control and the result of the other conditions are expressed as fold changes normalized to the empty control. Each condition was performed in technical duplicates and the experiments were performed in four independent biological replicates.

### Mopeia virus (MOPV) infections in the presence of SAMD9L

293T cells were seeded in 24-well plates at 0.2M cells per well. The following day, 1 µg of the SAMD9L plasmid was transfected with Lipofectamine™2000 (ThermoFisher Scientific), following the manufacturer’s instructions. In addition, 1 µg of a plasmid coding for mCherry was used as a transfection positive control and verified by fluorescence microscopy before 293T infection. For infection, 293T cells expressing mCherry or SAMD9L were infected with a recombinant Mopeia virus (MOPV_wt_) ^56^ at an MOI of 0.01. Inocula were prepared by diluting viruses in DMEM 2% FCS, 0.5% P/S. Cell supernatants were removed and viral inocula were added to corresponding wells. After a 1h incubation, wells were washed and reincubated for 48h. The cell supernatants of each condition were then collected and frozen at -80°C. Cells were collected and lysed in 2X Laemmli buffer and analyzed by western-blot to control for the expression of SAMD9L. For the titration of MOPV, Vero E6 cells were seeded in 12-well plates at 0.25M cells per well in DMEM 5% FCS, 0.5% P/S. The following day, cells were incubated with successive decimal dilutions of each sample supernatant for 1hr at 37°C and 5% CO_2_. After an hour, 1.5mL of a carboxymethylcellulose (CMC) solution was added to each well and plates were incubated at 37°C, 5% CO_2_ for seven days. CMC was then removed and cells were fixed in PBS 4% formaldehyde and permeabilized in PBS 0.5% Triton X-100. Focal Forming Units (FFU) were detected using rabbit primary antibodies specific to the MOPV Z protein (Agrobio) and secondary alkaline phosphatase-conjugated antibodies (Sigma-Aldrich Merck). Plates were revealed using an NBT/BCiP solution (ThermoFisher Scientific) and foci were counted to determine the virus titers. Viral titers were expressed as Focus-Forming Units per milliliter (FFU/mL).

### Host and viral sequence analyses

SAMD9 and SAMD9L orthologs from mammals were retrieved from public online databases using NCBI BLASTn with the human sequences of SAMD9 and SAMD9L as queries. SLFN sequences were retrieved using NCBI. HIV and lentivirus sequences were retrieved using the HIV sequence database http://www.hiv.lanl.gov/. Translation alignments were performed using Muscle ^57^. Geneious (Biomatters) and ESPript 3.0 ^58^ were used for sequence displays.

### Codon usage analysis

Codon usage preferences were measured using COUSIN (COdon Usage Similarity INdex) ^39^ through the online tool https://cousin.ird.fr/ with 18 492 transcripts from human ^39^, 223 from VACV, 4 from VSV, 4 from MOPV, 3 from MLV, 9 from HIV-1 LAI, 9 from HIV-1 pWITO, and a total of 43 337 transcripts from different HIV and SIV strains obtained through the HIV sequence database http://www.hiv.lanl.gov/.

### Other softwares and statistical analyses

Sequencing analyses and representations were performed in Geneious (Biomatters). Quantification of images (Western-blot analyses) were performed using ImageJ https://www.imagej.org. Graphic representations and statistics were performed using GraphPad Prism 9 (except some with Excel from Microsoft Office). In Figures, data are represented as mean ± SD. For Fig. 3, statistics were performed using the paired t-test. For Fig. 4 and Fig. S9, statistics were performed using the Mann–Whitney test.

## Data and reagents availability

All data and reagents are available upon reasonable request to the corresponding author.

## Acknowledgements

We thank Clarisse Berlioz-Torrent, Michael Emerman, Oliver Fregoso, Caroline Goujon, Molly OhAinle, and the LP2L members for helpful discussions on this project. We thank the Etablissement Francais du Sang for the leukopacks from blood donors and Xuan-Nhi Nguyen (CIRI) for some of the primary cell purifications. We thank Jerome Bourret, Philippe Paget-Bailly and Ignacio Bravo for discussion and support with the COUSIN analyses. We acknowledge the contribution of the SFR BioSciences (UAR3444/CNRS, US8/Inserm, ENS de Lyon, UCBL), PLATIM microscopy and ANIRA cytometry platforms, especially Jacques Brocard and Véronique Barateau for their help. We thank all the contributors of publicly available genome sequences and publicly available softwares. We thank Yan Xiang (UTHSCSA), Hirotaka Matsui (Kumamoto U), Yenan Bryceson (Karolinska Institutet), Xuan-Nhi Nguyen (CIRI), Sylvain Baize (CIRI) and the NIH AIDS reagent program for sharing reagents.

## Author Contributions

Conceptualization: AL, LE

Methodology: AL, FF, ER, MM, LE

Validation: AL, CD, CDMM, CL, LE

Formal analyses: AL, LE

Investigation : AL, CD, CDMM, CL, KN, MP, MW, FF, ER, LE

Resources : AL, AC, ER, MM, LG, LE

Writing – original draft: AL, LE

Writing – review and approved: all

Supervision: LE

Project administration: LE

Funding acquisition: LE

## Supplementary Information

### Supplementary Figures

**Figure S1.**
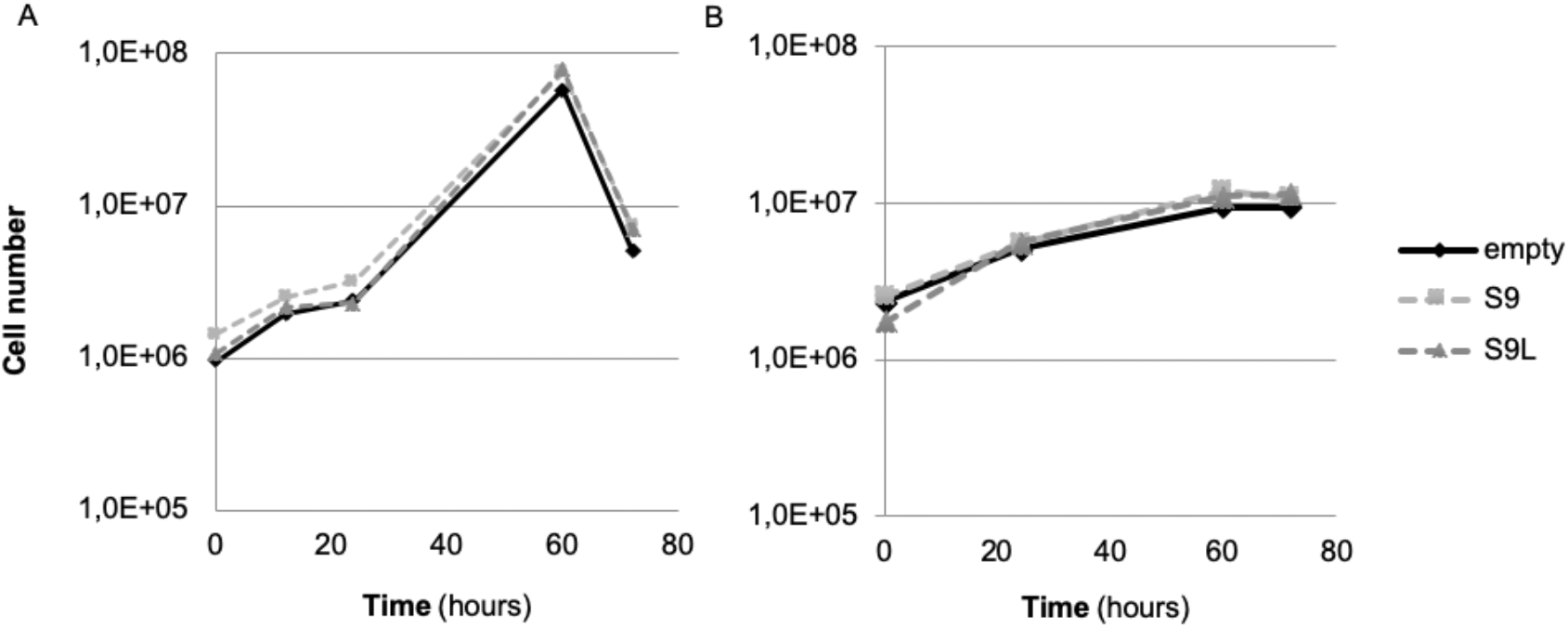
Overexpression of SAMD9 and SAMD9L in the conditions for the virus replication and infectivity assays do not impact cell proliferation. Panels A and B are from two independent experiments. Each condition was performed in technical duplicates.

**Figure S2.**
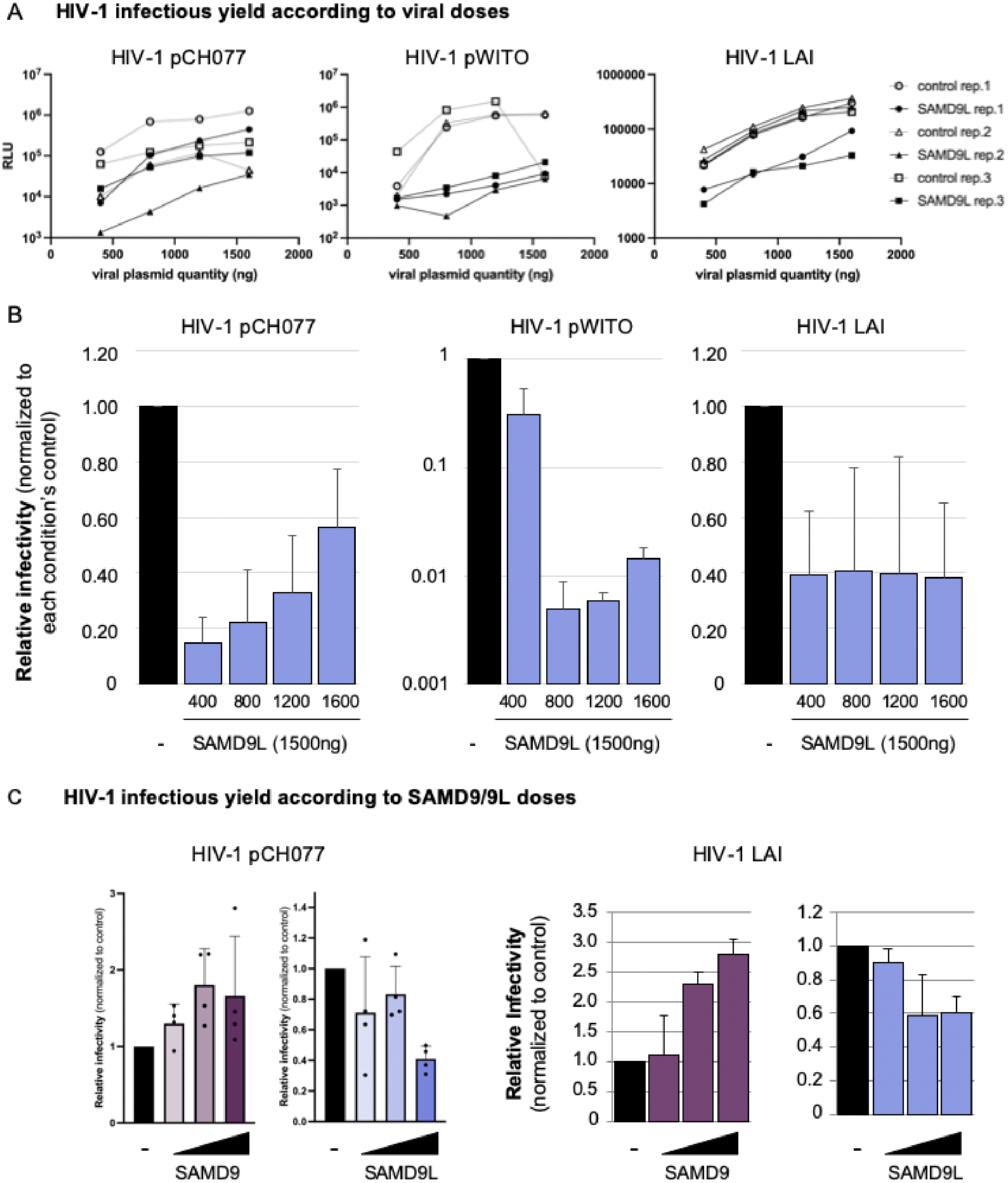
Identification of SAMD9L as a specific restriction factor of HIV-1 infectivity. A-B, HIV-1 infectious yield from +/-SAMD9L expressing cells according to the viral doses. Infectious yield of HIV-1 pCH077, pWITO, and LAI from supernatant of 293T cells co-tranfected with empty plasmid (control) or 1500 ng of SAMD9L plasmid and with 400, 800, 1200 or 1600 ng of viral-encoding plasmid. Virus infectivity was measured by luminescence (RLU) in the TZM-bl cell assay, where the luciferase reporter is under the LTR promoter (as in Fig. 1A). A, Raw RLU for the three independent biological replicates (Rep. 1-3). B, Relative infectivity: Mean of the experiments in A with a normalization to the empty control (in black). C, HIV-1 infectious yield from +/-SAMD9/9L expressing cells according to the SAMD9/9L doses. Relative infectivity of HIV-1 pCH077 and LAI from cells expressing SAMD9L or SAMD9 normalized to the empty control. 293T cells were co-transfected with empty plasmid (control) or 400, 1000 or 1500 ng of SAMD9L or SAMD9 plasmid, and with HIV-1 pCH077 or LAI plasmid.

**Figure S3.**
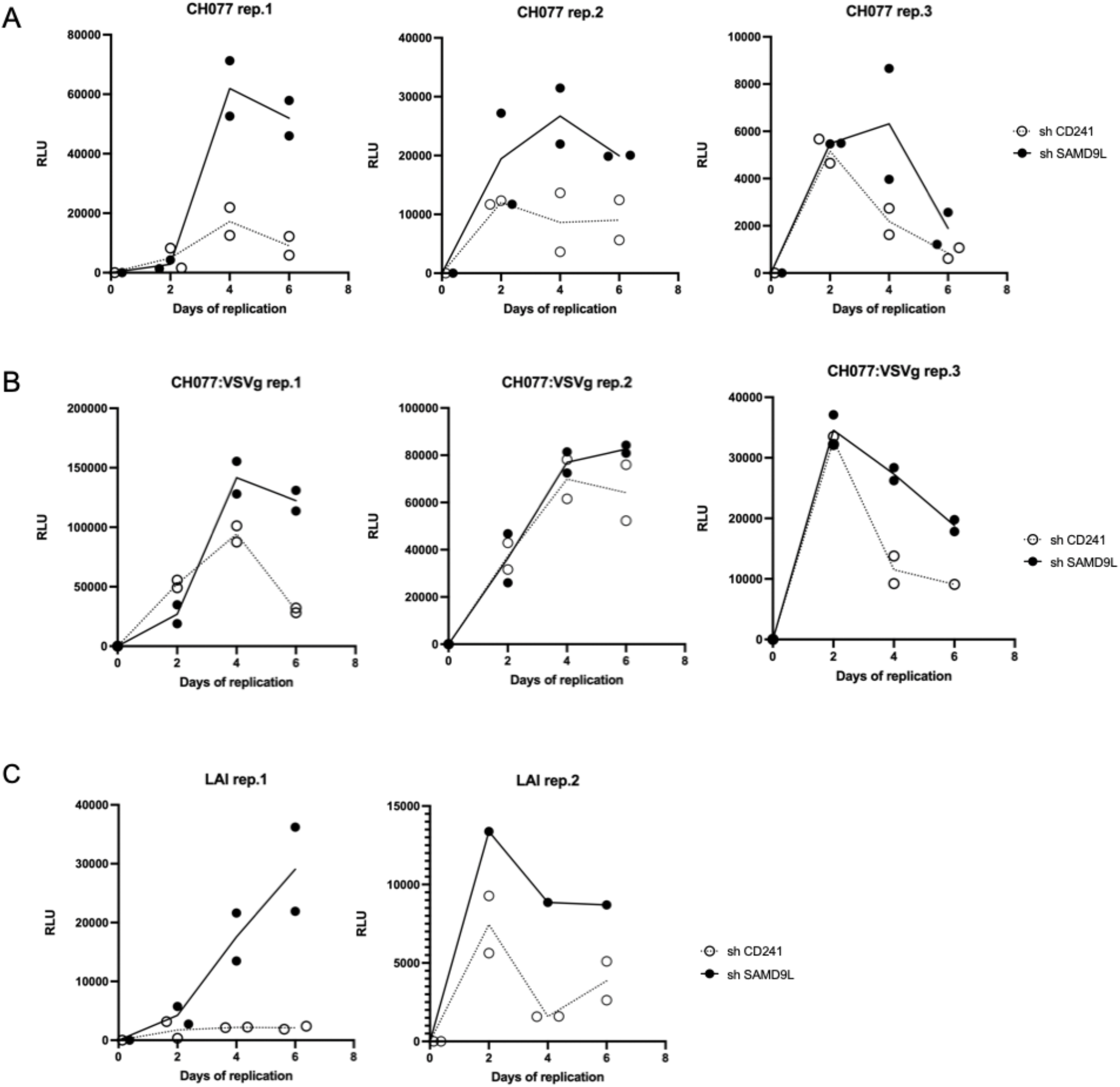
Endogenous interferon-stimulated SAMD9L restricts HIV-1 replication. HIV-1 T/F pCH077 (A), pCH077:VSVg (B) and pLAI (C) replication assay in shCD241 (control) or shSAMD9L HeLaP4P5 +IFNβ over 6 days of replication. Experimental setup shown in Fig 1D. Titration was performed using the TZM-bl assay. Each graph corresponds to an independent biological replicate (rep.1-3) with the technical duplicates for each condition. The Y-axis represents the raw RLU results. The first graph is also shown in Fig. 1F.

**Figure S4.**
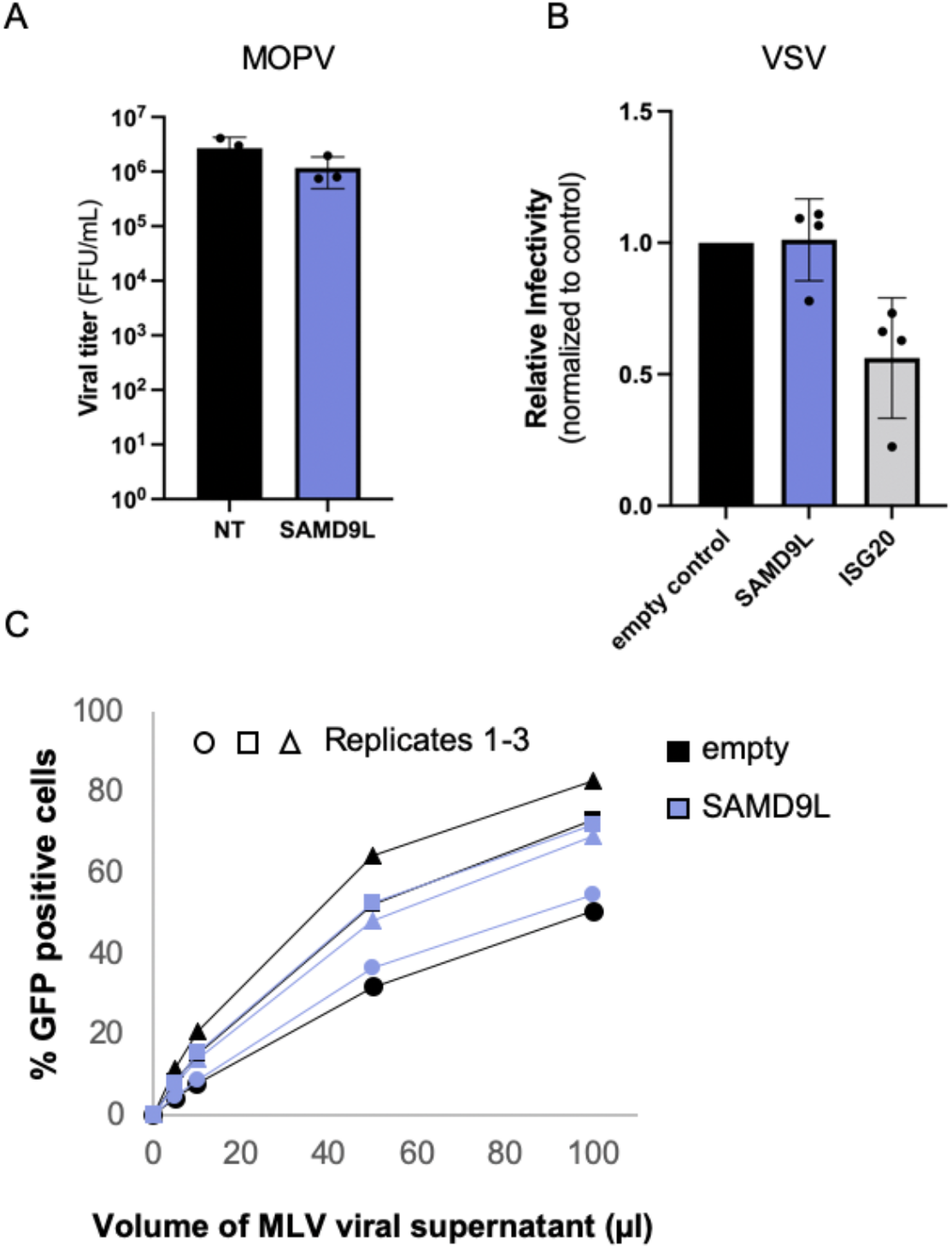
SAMD9L does not restrict the Arenavirus MOPV, the Rhabdovirus VSV, nor the Retrovirus MLV. A, MOPV viral titers retrieved from control cells (NT) or cells overexpressing SAMD9L. Titers were measured as Focal Forming Units (FFU). B, 293T cells transfected with an empty vector, SAMD9L or ISG20 (as positive restriction control) were infected 24h later with VSV:GFP replicative particles and supernatant was collected 16 hpi. The viral titer was measured in new cells by the % of GFP positive cells. Results are expressed as normalized to the empty control (condition in the absence of SAMD9L or ISG20). C, 293T cells were co-transfected with a SAMD9L or an empty plasmid, and with plasmids to encode for MLV:VSVg pseudoviruses: pTG5349 (MLV gagpol), PTG13077 (MLV LTR-GFP) and pMD2.G (VSVg). Viral supernatant was collected 48h later and 0, 5, 10, 50, 100 µl of supernatant was used to infect new target cells. Two days later, infectivity was measured by FACS (% of GFP positive cells) in the empty (in black) and SAMD9L (in blue) conditions.Results are shown for three independent biological replicates. Fig. 1H is the mean of these replicates.

**Figure S5.**
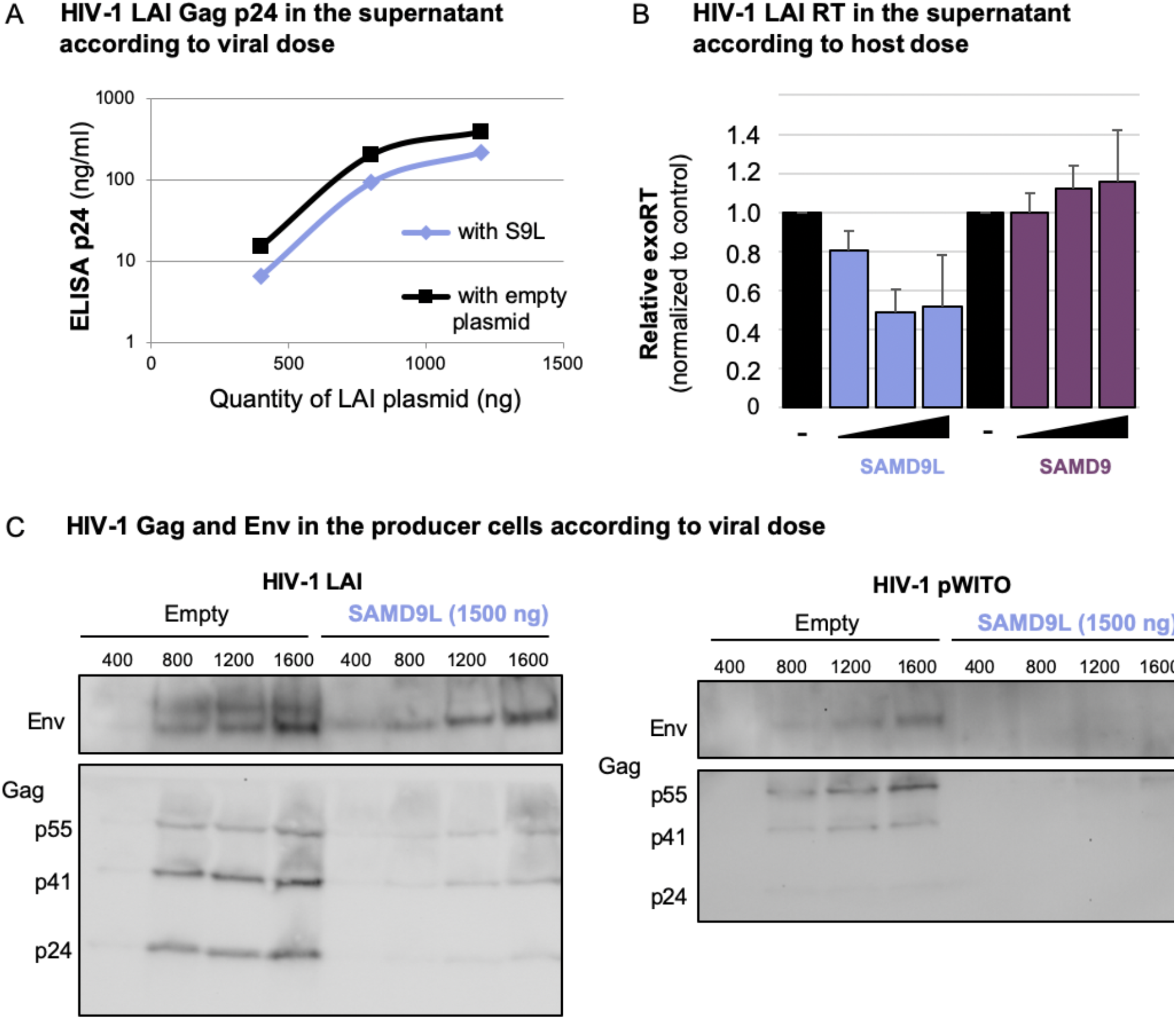
HIV-1 protein expression in the supernatant and in the producer cells is restricted by SAMD9L, and is not overcome by viral dose. A, Titration by ELISA p24 Gag of HIV-1 LAI in the supernatant of the producer cells overexpressing or not SAMD9L in the context of a dose of pLAI IMC plasmid (400, 800, 1200 ng). B, Titration by RT activity of HIV-1 LAI in the supernatant of the producer cells overexpressing or not SAMD9/9L in the context of a dose of SAMD9L or SAMD9 plasmid. A-B, Experiments were performed in three biological replicates, means of the replicates are shown here. C, Western-blot analyses of HIV-1 Gag and HIV-1 Env from lysates of the HIV-1 producer cells overexpressing or not SAMD9L in the context of HIV-1 pLAI (left) or pWITO dose (right) (IMC plasmids: 400, 800, 1200, 1600 ng).

**Figure S6.**
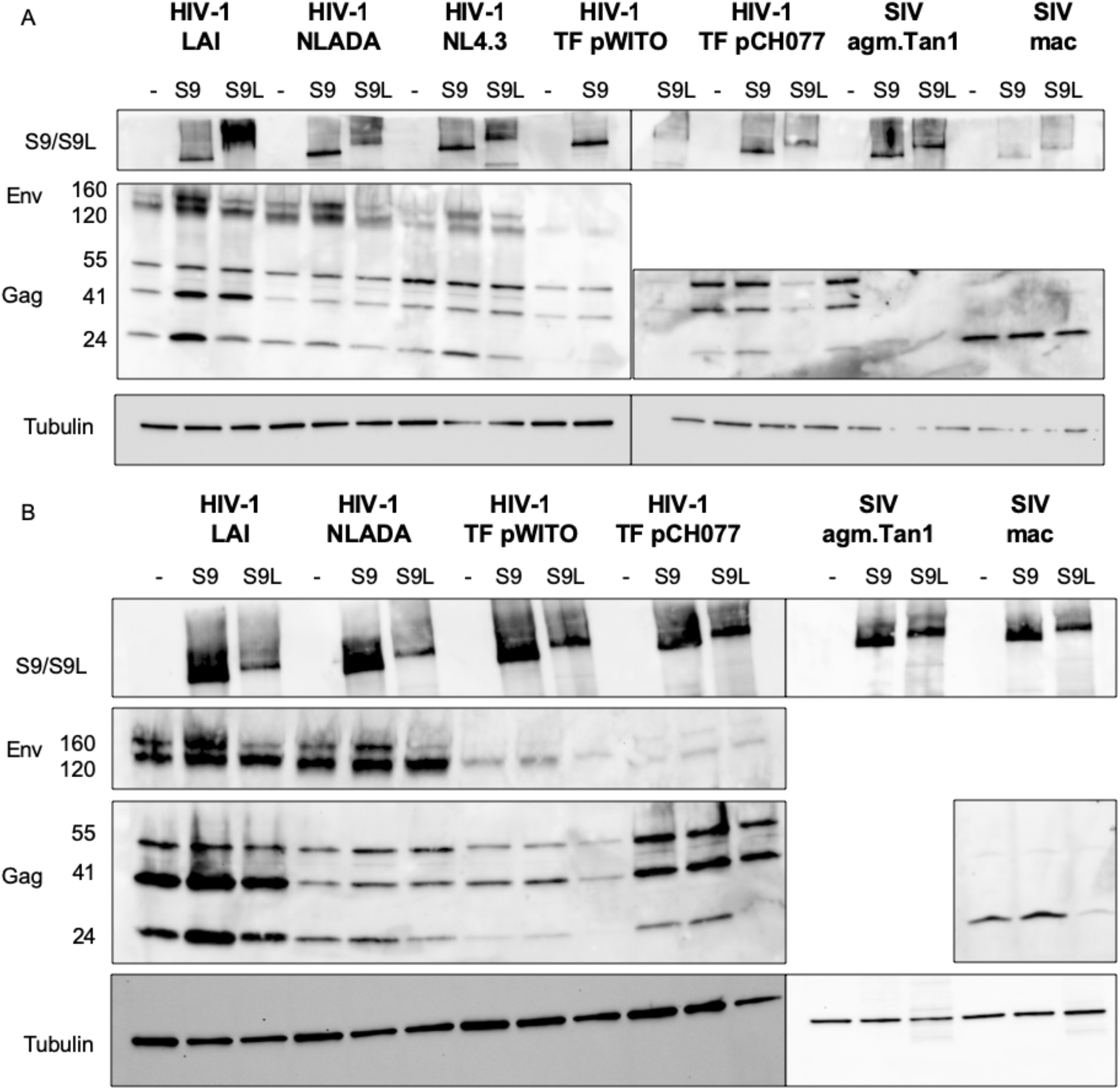
HIV-1 and SIV protein expression is inhibited by SAMD9L and enhanced by SAMD9. Western-blot analyses of SAMD9/9L, HIV-1/SIV Gag and Env, with Tubulin as loading control, from lysates of the viral producer cells in the context of overexpressed SAMD9/9L (1500 ng). A and B are from two independent biological replicates.

**Figure S7.**
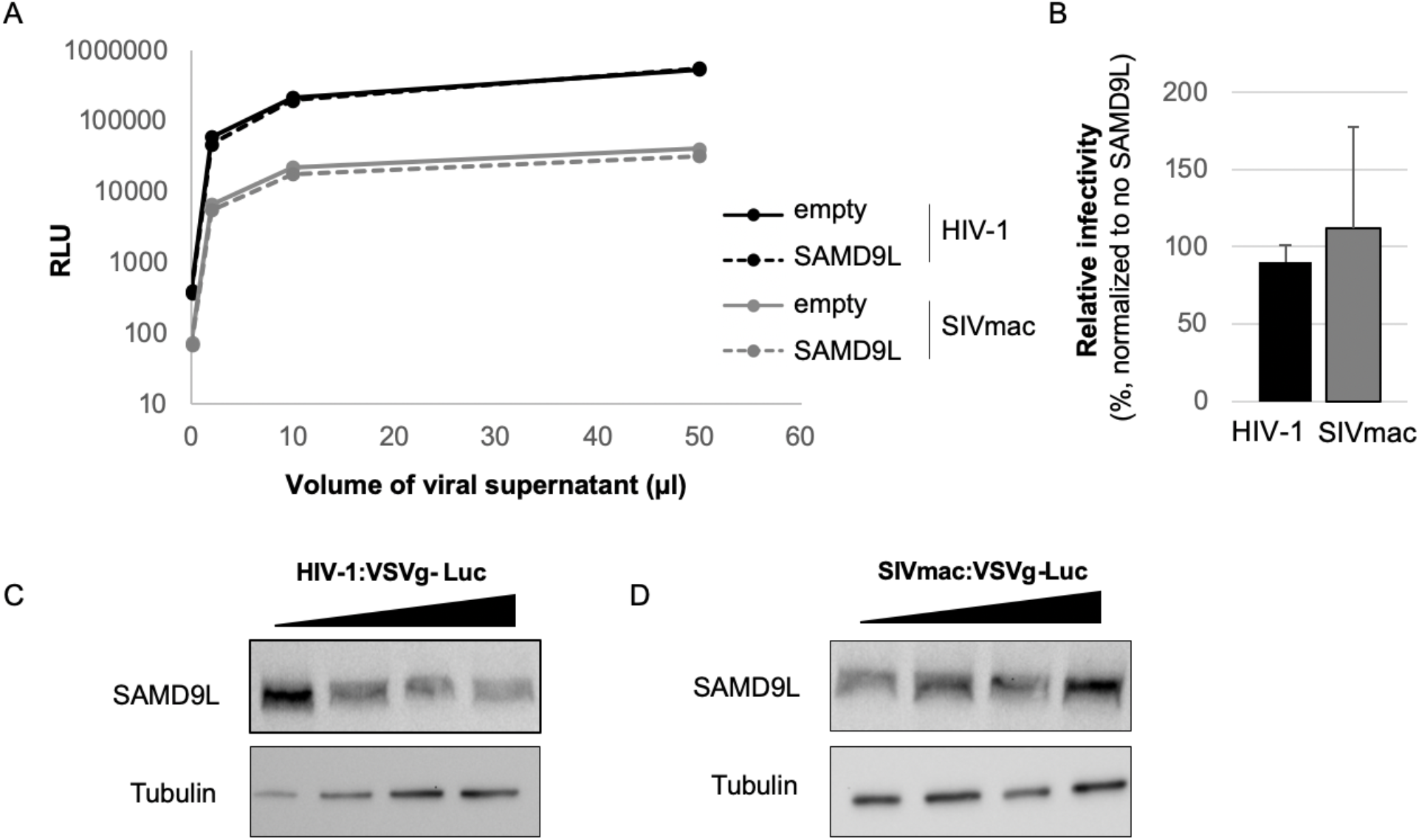
SAMD9L does not affect the early phases of primate lentivirus replication. A, Single-round lentiviral particles, HIV-1:VSVg-Luc and SIVmac:VSVg-Luc, were produced in 293T cells. Two days later, viral supernatant was collected and 1, 2, 10 or 50 µl was used to infect target cells overexpressing or not SAMD9L. Two days post-infection, cells were lysed and a luciferase assay was performed for titration. The Y-axis shows the relative light units (RLU) in the luciferase assay. The data represent the mean of three independent experiments. B, Corresponding relative infectivity of HIV-1:Luc and SIVmac:Luc (from the 10 µl condition in A), normalized to the no SAMD9L condition (100%). C-D, Expression of SAMD9L in target cells are not impacted by HIV-1 and SIVmac single-round infection. Western-blots showing SAMD9L cellular expression level two days post-infection with HIV-1:VSVg-Luc (C) or SIVmac:VSVg-Luc (D).Tubulin serves as a loading control.

**Figure S8.**
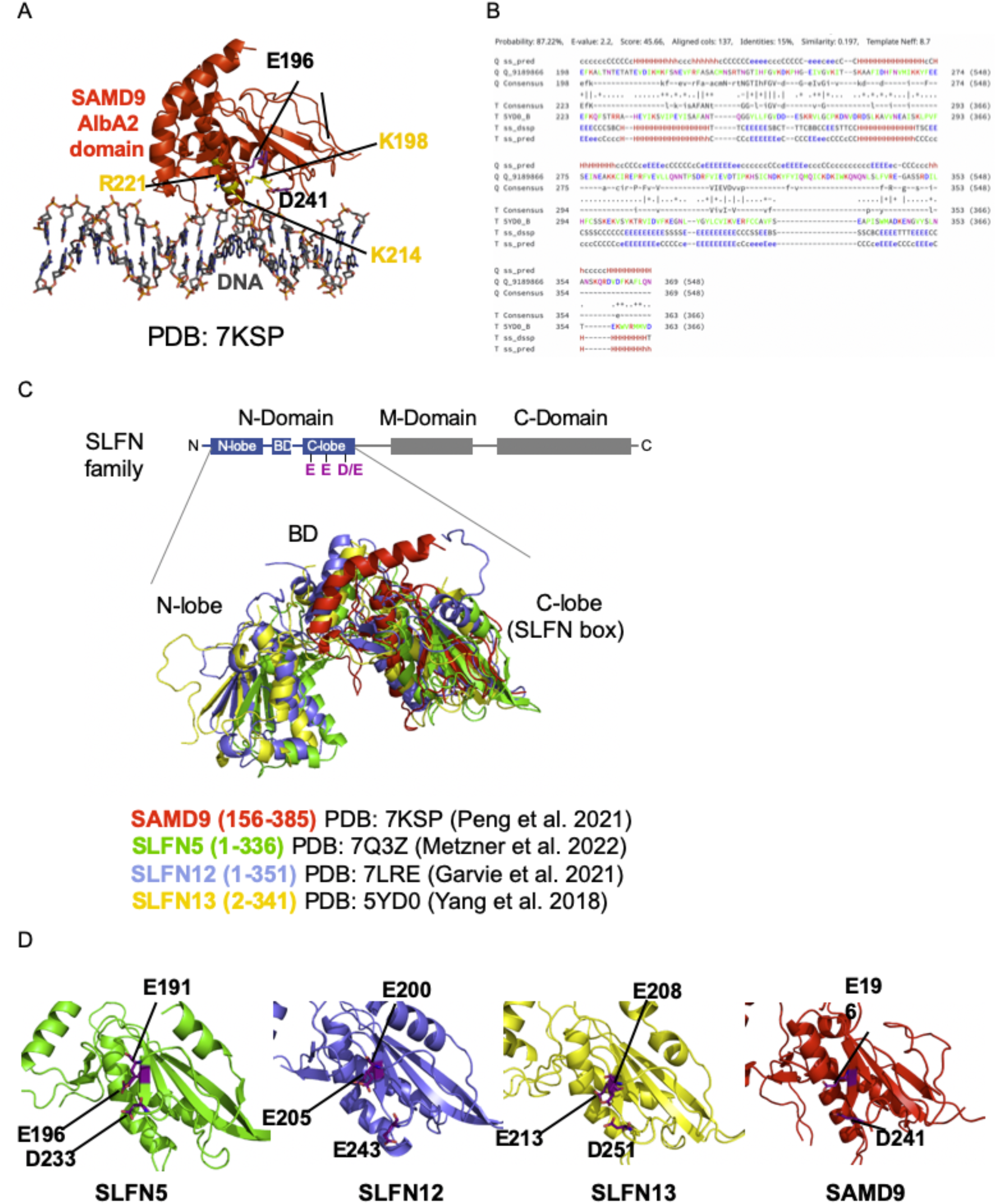
SAMD9 homology with SLFN proteins. A, SAMD9 crystal structure in complex with DNA (PDB 7KSP). Atoms of the residues forming the active site are represented with purple sticks (labeled in black), while those involved in DNA binding are in yellow. B, Snapshot of the alignment of rSLFN13 and SAMD9L from HHpred analysis results. C, Structural overlap of SLFN N-Domain including the SLFN-box of SLFN5, SLFN12, SLFN13 and SAMD9 AlbA2 domain. Residues forming the active site are indicated in purple below the schematic representation of SLFN family’s domains. D, Representation of the SLFN box, including the putative active site in purple, of SLFN5, SLFN12, SLFN13 and SAMD9. Coordinates are labeled in black.

**Figure S9.**
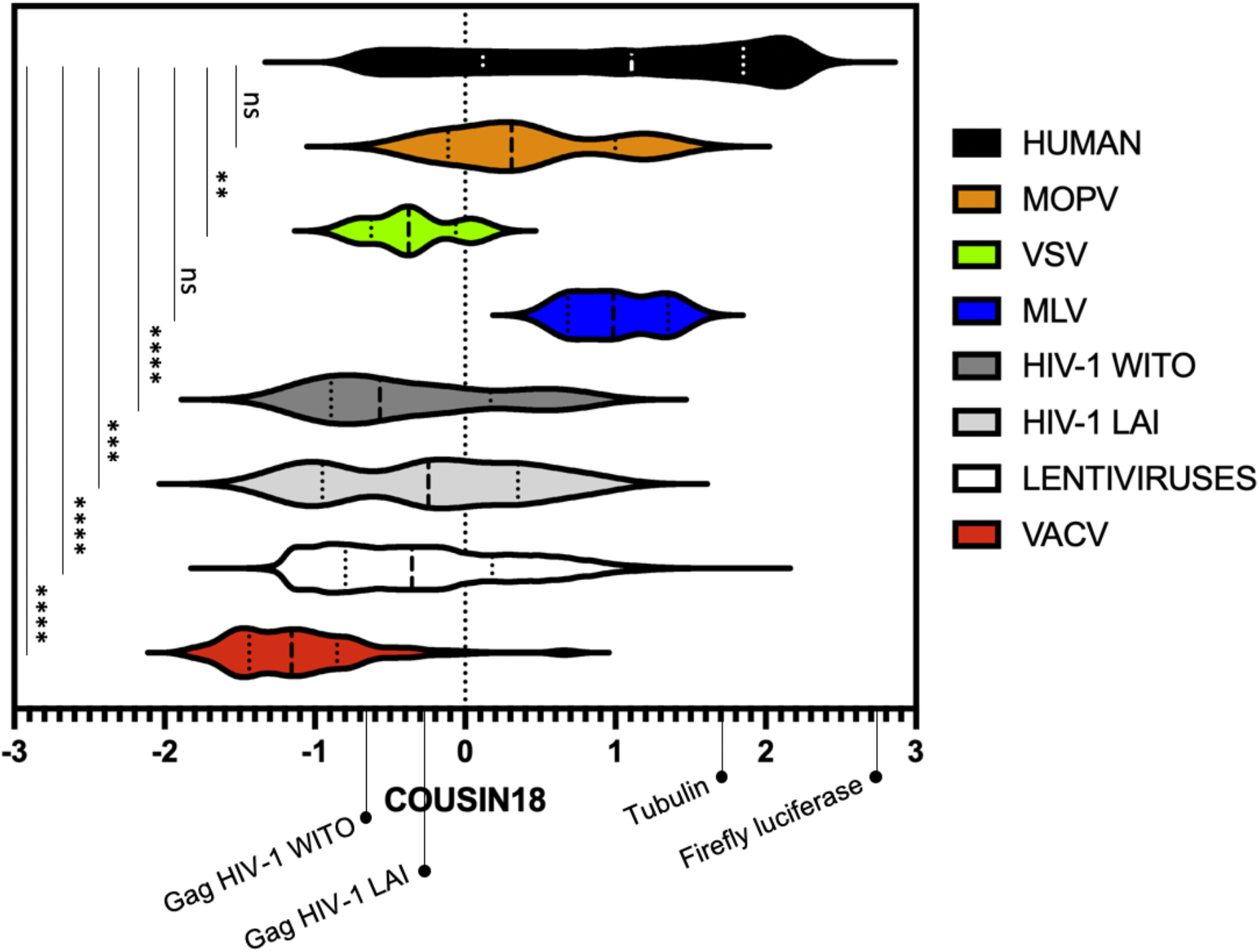
Codon usage of viral transcripts from Arenavirus MOPV, Rhabdovirus VSV, Gammaretrovirus MLV, Poxvirus VACV and Lentiviruses as compared to the human ones. Genome-wide codon usage analyses of human and viral transcripts using the COUSIN18 score for Human (18,492 gene transcripts from Bourret et al. 2019), MOPV (4), VSV (4), MLV (3), HIV-1 pWITO (9), HIV-1 LAI (9), Lentiviruses HIVs/SIVs (43,337), VACV (223). COUSIN18 scores of selected transcripts (Gag from HIV-1 pWITO and pLAI, Tubulin, and Firefly Luciferase) are further highlighted. ****, ***, **, and ns are p-values < 0.0001, < 0.001, < 0.01, and >0.05, respectively.

### Supplementary Tables

**Table S1.**
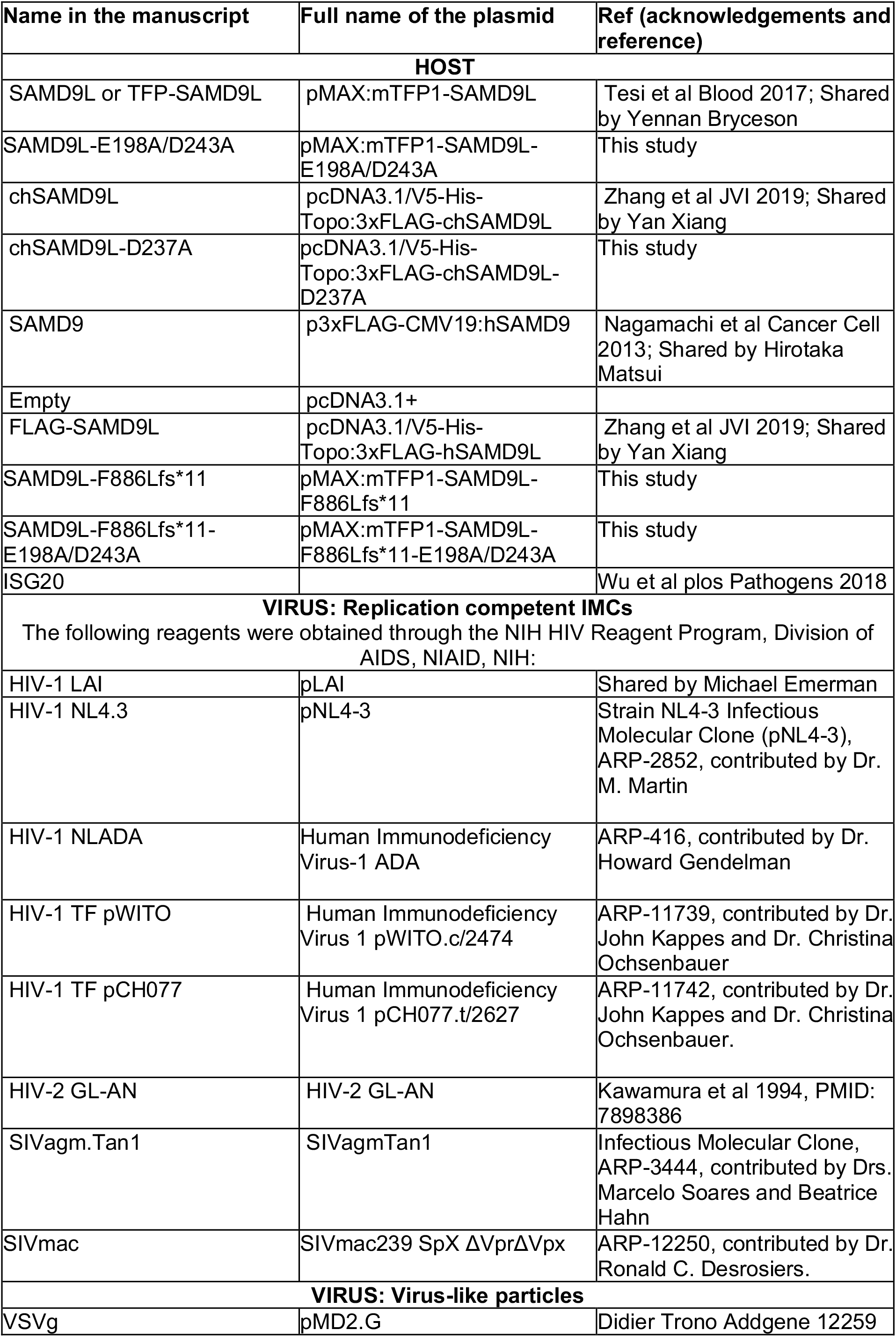
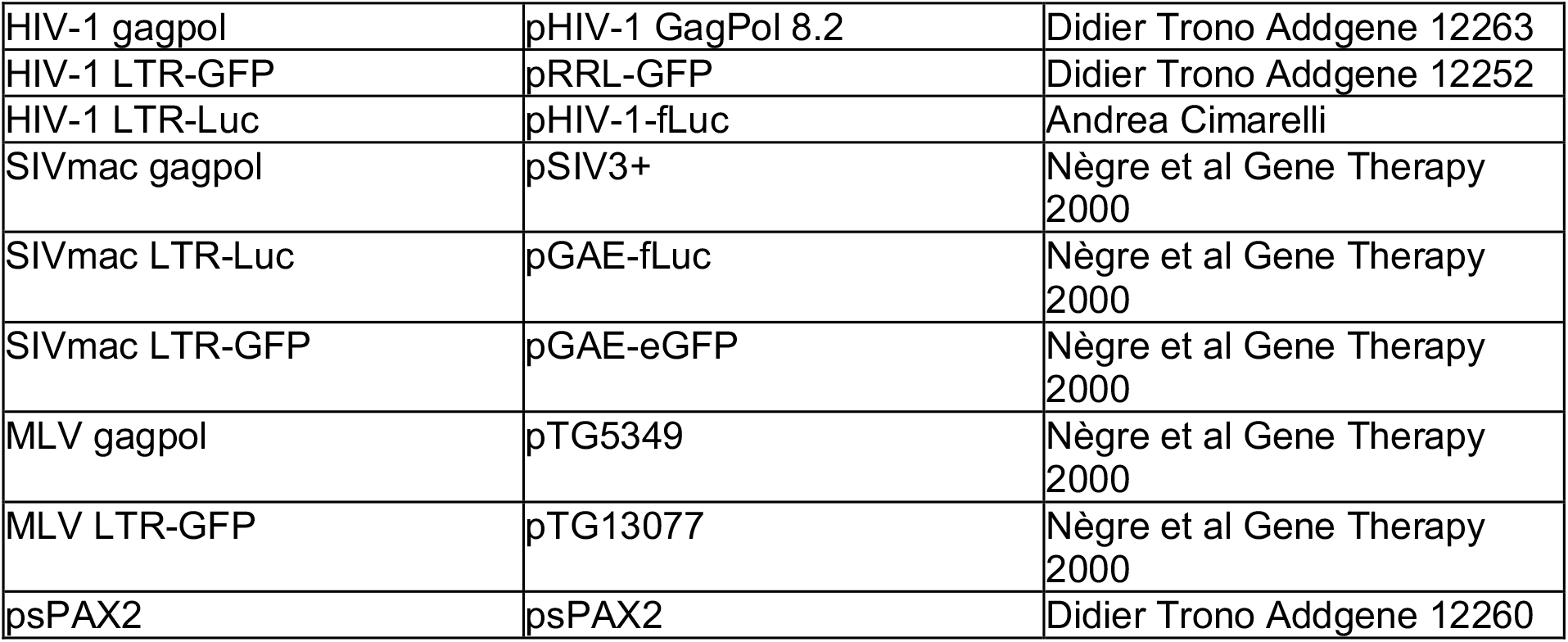
Plasmids used in the study.

**Table S2.**
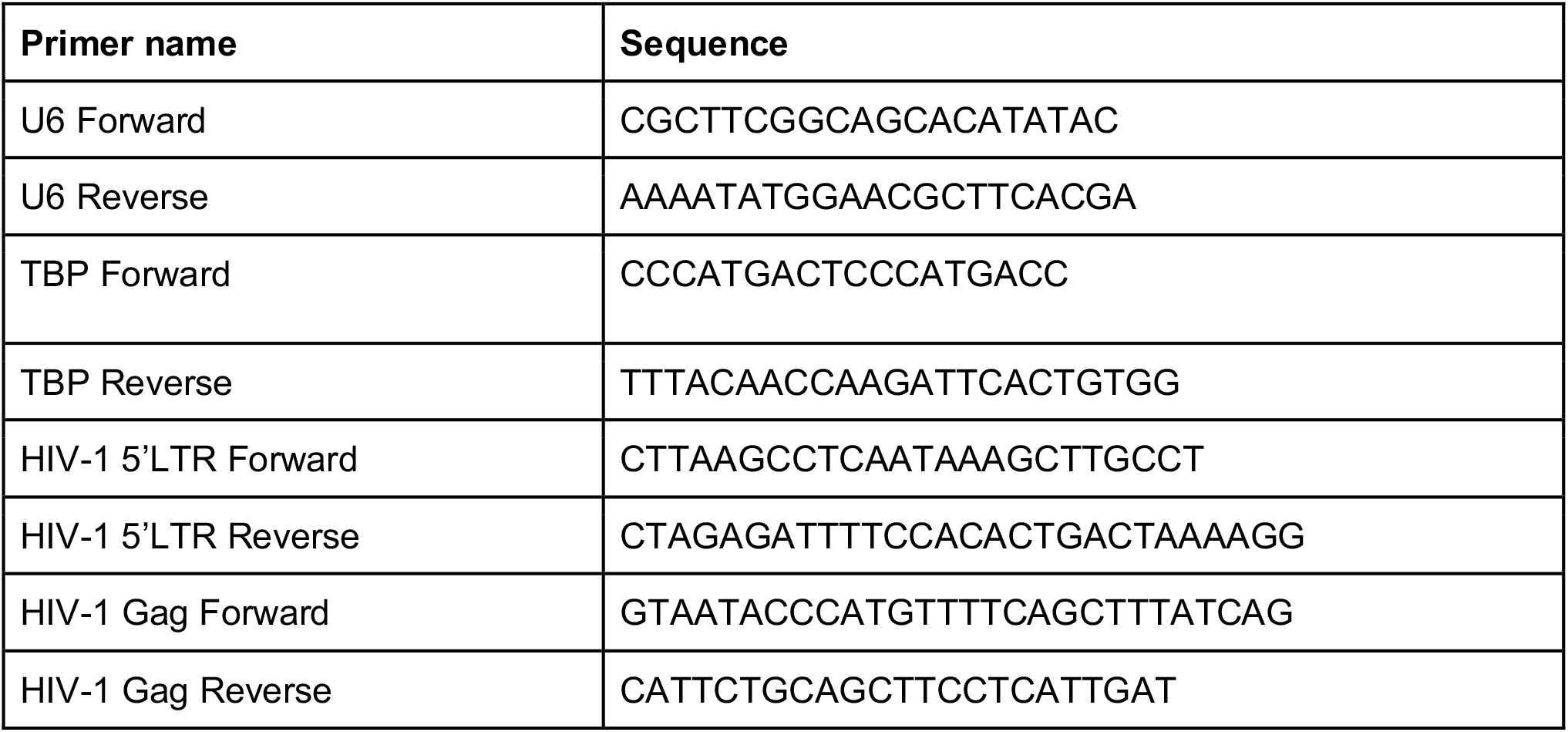
Primers for qPCR used in the study.

